# A novel role for cystathionine γ lyase in the control of p53: impact on endothelial senescence and metabolic reprograming

**DOI:** 10.1101/2022.09.05.506654

**Authors:** Jiong Hu, Matthias S. Leisegang, Mario Looso, Gabrijela Dumbovic, Janina Wittig, Maria-Kyriaki Drekolia, Stefan Guenther, David John, Mauro Siragusa, Sven Zukunft, James Oo, Ilka Wittig, Susanne Hille, Andreas Weigert, Stefan Knapp, Ralf P. Brandes, Oliver J. Müller, Andreas Papapetropoulos, Fragiska Sigala, Gergana Dobreva, Ingrid Fleming, Sofia-Iris Bibli

**Author notes:** **First author**: Jiong Hu, PhD. **Correspondence:** Sofia Iris Bibli PhD and Ingrid Fleming PhD. Goethe University Frankfurt, Institute for Vascular Signalling, Centre for Molecular Medicine, Theodor Stern Kai 7, 60596 Frankfurt am Main, Germany.

## Abstract

**Aims:** Advanced age is unequivocally linked to the development of cardiovascular disease, however, the mechanisms leading to loss of endothelial cell regenerative capacity during aging remain poorly understood. Here we aimed to investigate novel mechanisms involved in endothelial cell senescence, that impact on endothelial cell transcription and the vascular repair response upon injury

**Methods and results:** RNA sequencing of a unique collection of native endothelial cells from young and aged individuals, showed that aging (20 vs. 80 years) is characterized by p53- mediated reprogramming to promote the expression of senescence-associate genes. Molecular analysis revelead that p53 accumulated and acetylated in the nucleus of aged human endothelial cells to suppress glycolysis. Metabolic flux analysis identified an associated reduction in glucose uptake and ATP availability that inhibited the assembly of the telomerase complex, which was essential for proliferation. Nuclear translocation of p53 in aged endothelial cells was attributed to the loss of the vasoprotective enzyme, cystathionine γ-lyase (CSE), which physically anchored p53 in the cytosol. In mice, loss of endothelial cell CSE activated p53 and arrested vascular repair upon injury, while the AAV9 mediated re-expression of an active CSE mutant retained p53 in the cytosol, maintained endothelial glucose metabolism and proliferation, and prevented endothelial cell senescence. Adenoviral overexpression of CSE in human native aged endothelial cells maintained low p53 activity and re-activated telomerase to revert endothelial cell senescence.

**Conclusion:** Our data identified the interaction between CSE and p53 as a promising target to preserve vascular regeneration during aging.

**Key Question:** To identify the mechanisms that regulate endothelial cell senescence under native conditions and their impact on vascular repair in aging.

**Key Finding:** Lack of a physical interaction between CSE and p53 metabolically reprogrammes endothelial cells to reduce telomerase activity and halt endothelial cell regeneration.

**Take home message:** Interventions to increase CSE expression represent a novel therapy against p53-induced endothelial cell cycle arrest and senescense

**Translational perspective:** Endothelial rejuvenation strategies could serve as promising therapies against age-related cardiovascular diseases. By investigating human native endothelial cells from young and aged individuals, we identified that the age-related nuclear accumulation of p53 reprograms endothelial cell metabolism, regulates telomerase activity and inhibits endothelial cell regeneration. Nuclear localization of p53 resulted from a loss of its interaction with the cysteine catabolizing enzyme cystathionine γ-lyase in the cytoplasm. Enhancing the physical interaction of p53 with CSE by gene therapy could revert endothelial cell senescence and activate endothelial reparative responses.

## Introduction

Advanced age is the dominant unmodifiable risk factor for the development of cardiovascular disease.^1,2^ Large landmark studies have demonstrated that changes associated with vascular aging; such as chronic low-grade inflammation, oxidative stress and endothelial dysfunction, are positively correlated with telomere attrition, cellular senescence, dysregulated nutrient sensing and stem cell exhaustion.^3,4^

A surprisingly large number of aging pathologies have been linked to senescence i.e. the irreversible arrest of cell proliferation that occurs when cells experience prolonged stress.^5^ To-date, senescence has been associated with two major tumor suppressor pathways i.e., p53/p21 and p16INK4a/pRB.^6^ No known physiological stimuli can stimulate senescent cells to re-enter the cell cycle but molecular manipulations, such as the sequential inactivation of certain tumor suppressor genes, can induce senescent cells to proliferate^6^. Notably, the elimination of senescent cells (senolysis) was recently reported to improve normal and pathological changes associated with aging in mice^7,8^ and the recent development of endothelial cell-specific senolytic targeting seno-antigens could be a promising strategy for new therapies to diminish pathological phenotypes associated with aging.^9^

Much of our knowledge regarding the effects of aging on the vasculature has been derived from a combination of cell culture studies; using multi-passaged cells or immortalized cells, and investigations in rodents.^10–12^ Although recent study reported the accumulation of senescent cells and the differential regulation of the Wnt pathway in the supernatant from explanted mammary arteries from older human subjects,^13^ there is a real lack of detailed information about the native aged human endothelium.

In this study we made use of a unique collection of human native endothelial cells from young and old subjects to investigate novel mechanisms involved in endothelial cell senescence, that impact on endothelial cell transcription and the vascular repair response following upon injury.

## Methods

Methodological details are provided in the Supplemetary Online Material.

## Results

### p53 Lys120 acetylation is a key regulator of endothelial cell senescence

Mesenteric arteries were obtained from young (20±3.4 years) and aged (80±2.3 years) individuals during interventions for conditions not related to cardiovascular disease. Native endothelial cells (CD144^+^ cells) were isolated from the arteries, FACS sorted within 45 minutes, and frozen after a short recovery period (20 minutes) in Hank’s buffer supplemented with 5% orthologous serum. Endothelial cells from aged donors were characterized as senescent as they demonstrated a significant reduction in telomere length (**Figure 1A**), and an increase in the activity of senescence-associated β-galactosidase (SAβG) compared to cells from young donors (**Figure 1B**). To assess transcriptional changes and identify upstream regulatory factors driving endothelial cell aging we performed RNA sequencing. Transcriptome (**Figure 1C**) and gene set enrichment analyses (**Figure 1D**) revealed the over-representation of pathways involved in *cell cycle control* and *p53-mediated transcription* in aged endothelial cells. Indeed, several p53 target genes were upregulated in aged endothelial cells, including p21 (CDKN1A),^14^ and the E3 ubiquitin ligase mouse double minute 2 or MDM2 (**Figure 1E**). The transcription factor p53 and the cyclin-dependent kinases, p16 and p21, are reported to drive senescence^15,16^ and mRNA encoding all three transcripts were increased in aged human endothelial cells (**Figure 1F**). When we investigated chromatin accessibility to the p53 binding motif (ATAC sequencing) in the young and aged endothelium no global changes were evident (**Figure 1G**). However, more p53 was detected in the nucleus of the endothelial cells from aged individuals, indicating increased p53 activity (**Figure 1H**). Non-acetylated nuclear p53 is inactive,^17^ but the acetylation of p53 on Lys120 was increased in nuclei from aged endothelial cells (**Figure 1I**). Next, endothelial cell senescence was induced *in vitro* in cells from young individuals using a multi-duplication approach (passaging). To address the impact of altering p53 expression on endothelial senecence, p53 was deleted (using a guide RNA lentiviral CRISPR approach), followed by lentiviral mediated overexpression of a wild-type p53 or a non-acetylatable p53-K120R mutant and cell proliferation was compared with cells infected with a non-targeted control lenti-CRISPR (**Figure 1J**). After 10 passages, replication was poor but comparable in cells treated with the non-targeted control or with the combination of p53 knockdown and overexpression of the wild-type p53. Cell proliferation was, however, significantly improved in cells lacking p53, as well as in cells expressing the non-acetylatable p53-K120R mutant (**Figure 1K**). Comparable effects were also observed on SAβG expression (**Figure 1L**). Taken together these data indicate that senescence of native human endothelial cells is linked to an increase in nuclear p53 levels and its acetylation on K120, which ensures its activation and impacts on endothelial proliferation.

**Figure 1.**
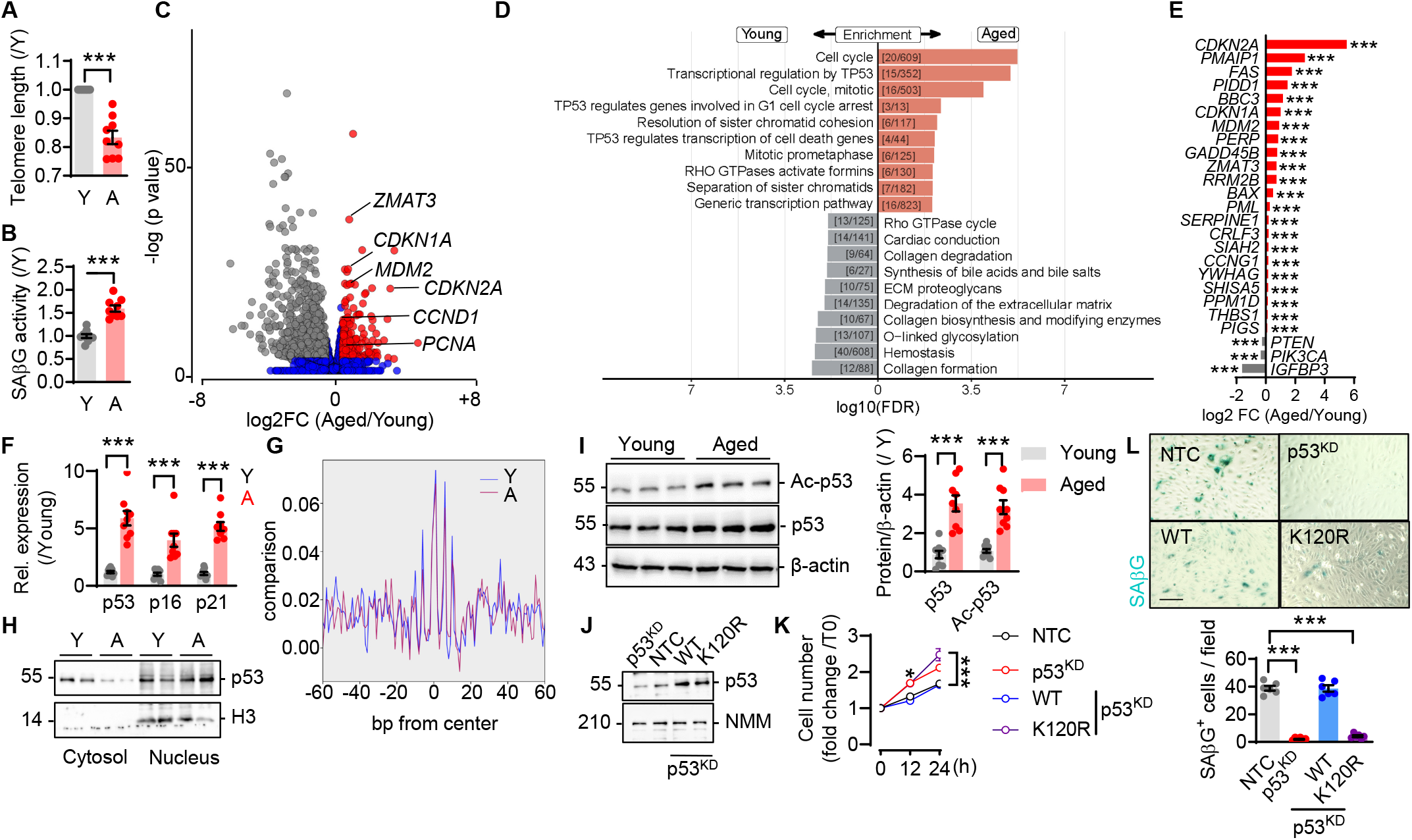
Acetylation of p53 on K120 regulates endothelial cell senescence and proliferation. Native mesenteric artery endothelial cells were isolated from young (Y) and aged (A) individuals; n=6-9/group with each sample a pool of 5 different human arteries. (**A**) Telomere length. (**B**) Senescence-associated β-galactosidase (SAβG) activity. (**C**) Volcano plot showing fold change (FC) in RNA transcripts (mRNA sequencing). (**D**) Gene set enrichment analysis (Reactome) using KOBAS. Top 10 gene sets or pathways enriched for up- or down-regulated genes (FDR<0.2) in samples as in panel C. Red: genes significantly upregulated in aged, grey: genes significant downregulated in aged. (**E**) Ratio of transcript expression in cells as in panel C for KEGG pathway 04115-enriched p53 targets. FC: fold change. (**F**) p53, p16 and p21 mRNA levels. (**G**) Overall chromatin accessibility at all p53 motif sites. (**H**) Representative immunoblot showing p53 expression in the cytosol and nucleus. (**I**) Expression of p53 and its acetylation (Ac) on K120 in whole cell lysates. (**J-L**) Endothelial cells from young individuals (passage 1) were treated with a non targeting control (NTC) or lentiCRISPR viruses targeting p53 (p53KD). Some p53KD cells were simultaneously transduced with lentiviruses to overexpress either wild-type p53 (WT) or a non-acetylatable mutant (p53K120R). Cells were then cultured for additional 9 passages. (J) Representative immunoblot of p53. (K) Fold change in cells cultured for up to 24 hours (at passage 10). (L) SAβG staining in endothelial cells from young individuals treated as in panel J. Bar = 50 μm. J-L, n=3 cell isolates in 2 independent experiments. ***P<0.001. Student’s t-test (A, B, E, F, I), two way ANOVA (Tukey’s, K, L).

### Impact of p53 on metabolic reprogramming and endothelial cell proliferation

p53 has been proposed to regulate cellular metabolism, in particular glycolysis,^18,19^ in part through the negative regulation of the glucose transporter Glut1 (also known as SLC2A1).^20^ As the metabolic consequences of aging had not been studied in native human endothelial cells, we used targeted metabolomics to assess changes in glycolysis, amino acid catabolism, TCA cycle and the pentose phosphate pathway. There were marked differences in the metabolic profiles of the endothelial cells from young and aged individuals, as indicated by orthogonal partial least square discriminant analysis (OPLS-DA) (**Figure 2A**, see Supplementary material online, **Figure S1A**). S-plots generated after multivariate OPLS-DA modeling identified 7 intracellular metabolites that were enriched in, and 10 metabolites depleted from, the aged human endothelium (**Figure 2B**, Supplementary material online, **Table S1**). Pathway impact and enrichment analysis of the most altered metabolites revealed differences in arginine, proline, cysteine and methionine metabolism as well as in glycolysis (see Supplementary material online, **Figure S1A-S1C**). Aged endothelial cells contained less glucose, glucose-6-phosphate and pyruvate than cells from younger individuals (**Figure 2C**), indicating that aging reduced endothelial cell glycolytic activity.

**Figure 2.**
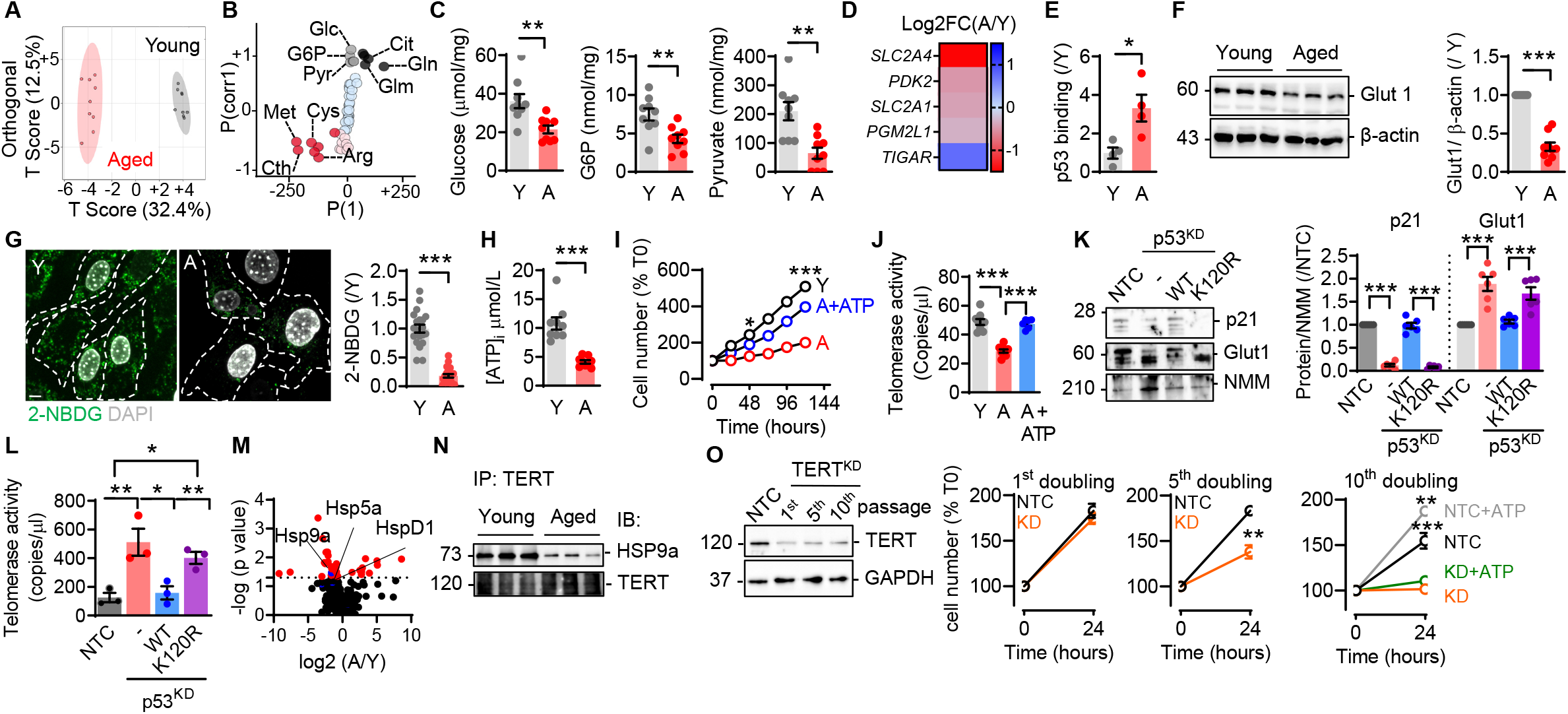
p53 induced metabolic reprogramming reduces ATP and impairs telomerase activity and endothelial proliferation in aging. Native mesenteric artery endothelial cells were isolated from young (Y) and aged (A) individuals; n=6-9/group with each sample a pool of 5 different human arteries. (**A**) Orthogonal partial least square discriminant analysis (OPLS-DA) 2D score plot of the metabolic profile. (**B**) OPLS-DA S-plot of the most abundant metabolites contributing to the T score difference presented in panel A. Annotations highlight metabolites deprived (black dots) or accumulated in aged endothelial cells (red dots). Glc: glucose; G6P: glucose-6 phosphate; Cit: citrulline; Pyr: pyruvate; Gln: glutamine; Glm: glutamate; Cys: cysteine; Met: methionine; Cth: cystathionine; Arg: arginine. (**C**) Intracellular levels of glucose, glucose-6 phosphate (G6P) and pyruvate. (**D**) Heatmap showing p53 related transcripts as log2 fold change (FC), RNA sequencing. (**E**) p53 binding to the Glut 1 promoter detected by ChlP- qPCR. (**F**) Glut 1 protein expression; n=4/group (each sample is a pool of 5 independent isolations). (**G**) 2-NBDG uptake. Bar = 2 μm. (**H**) Intracellular concentration of ATP. (**I**) Precentage of cell number from cells treated with solvent or ATP-polyamine-biotin every 24 hours. (**J**) Telomerase activity in cells as in panel I. (**K-L**) Endothelial cells from young donors were treated (in passage 1) with a non targeting control (NTC) or a p53 (p53KD) lentiCRISPR viruses. Some p53KD cells were left untreated (-) others were simultaneously transduced with lentiviruses overexpressing a wild-type p53 (WT) or a non-acetylatable mutant (K120R). Cells were then cultured for additional 9 passages. (K) Representative immunoblots and quantification of p21 and Glut1 and (L) telomerase activity. (**M**) Telomerase complex identified by LC/MS-MS. (**N**) Immunoprecipitation of human telomerase reverse transcriptase (TERT) and immunoblotting for TERT and HSP9a. (**O**) Human native young endothelial cells were treated with lentiCRISPR viruses carrying a non targeting control gRNA (NTC) or gRNA directed against TERT (KD). Representative western blot for TERT and effects on percentage of cell number for up to 10 passages. On passage 10, cells were treated with solvent or ATP-polyamine-biotin for 24 hours. K, L, O, n=3 per group, 2 independent experiments. *P<0.05, **P<0.01, ***P<0.001. Student’s t- test (C, E, F, G, H), one-way ANOVA (Bonferroni, J, K, L, O) and two-way ANOVA (Tukey’s, I, O).

In cancer cells, p53 has been shown to induce the expression of fructose-2,6-bisphosphatase (also known as TP53-induced glycolysis and apoptosis regulator or TIGAR), to inhibit phosphoglycerate mutase (PGM) and repress Glut1 and Glut4 expression.^19^ Consistent with the increase in p53 activity in aged endothelial cells, they expressed less Glut1 and Glut4 but higher levels of TIGAR than cells from younger donors (**Figure 2D**). By performing p53 ChIP followed by qPCR, we were able to show that the (repressive) binding of p53 to the Glut1 promoter was increased in endothelial cells from aged individuals (**Figure 2E**) and resulted in a decrease in Glut1 protein levels (**Figure 2F**). The expression of other glycolytic enzymes was comparable in the young and aged endothelium (see Supplementary material online, **Figure S1D**). Changes in Glut1 expression were correlated with reduced glucose import in aged cells as evidenced by impaired uptake of 2-NBDG, which was readily imported into endothelial cells isolated from young individuals (**Figure 2G**). Impaired glycolysis in aged endothelial cells also resulted in a marked reduction in intracellular ATP levels (**Figure 2H**). Similar changes in metabolism were also evident in native aortic endothelial cells from 12 versus 1 month old wild-type mice (see Supplementary material online, **Figure S2A-S2D**).

### Impact of p53-induced metabolic reprogramming on telomerase activity

To determine whether altered glucose metabolism could be directly linked with the aged phenotype, endothelial cells from young individuals were treated with siRNAs directed against Glut1. Decreasing Glut1 by 60% (see Supplementary material online, **Figure S3A**), resulted in a positive SAβG signal after 4 additional passages (see Supplementary material online, **Figure S3B**), as well as increased expression of p21 and p16 (see Supplementary material online, **Figure S3C**), versus cells treated with a control siRNA. Thus, interefering with glucose uptake accelerated the onset of endothelial cell senescence. The impact of metabolism on the development of senescence was further highlighted by treating aged cells with a cell-permeable ATP analogue, which significantly increased cell proliferation (**Figure 2I**). Consistent with the changes in telomere length (see Figure 1A), telomerase activity; as determined using a droplet digital telomere repeat amplification protocol (ddTRAP), was reduced in endothelial cells from aged individuals but was increased after addition of the ATP analogue (**Figure 2J**). Similarly, in multi-passaged endothelial cells, reduced telomerase activity was improved following the knockdown of p53. Simultaneous expression of the wild-type p53 but not the K120R mutant, abrogated the effect of p53-deletion on telomerase activity (**Figure 2L**). The downregulation of p53 also decreased p21 expression but increased Glut1 levels in multi-passaged endothelial cells, effects that were reversed by the overexpression of wild-type p53 but not the p53-K120R mutant (**Figure 2K**). Glut1 silencing also resulted in reduced telomerase activity (see Supplementary material online, **Figure S3D**), indicating that glucose metabolism indeed acts as a regulator of telomerase.

The assembly of the active telomerase complex is an energy requiring process (Lue and Wang 1995) that involves the recruitment of ATP-dependent chaperones.^21^ To determine whether endothelial cell aging had an impact on the telomerase complex, the catalytic subunit of the telomerase reverse transcriptase (TERT) was immunoprecipitated from young and aged endothelial cells and its interaction partners were assessed by mass spectrometry (Supplementary material online, **Table S2**). This approach revealed pronounced changes in TERT-associated proteins with aging (**Figure 2M**). In endothelial cells from young individuals, 5 chaperone proteins interacted with TERT, including its previously decribed interacting partner HSP90.^22^ The chaperone-TERT interactione was, however, not detectable in aged endothelial cells. One of the TERT-associated proteins was the ATP-dependent chaperone; HSP9a (also known as mortalin), and its association with TERT in endothelial cells from young individiulas was confirmed by immunoprecipitation, as was its decreased association with TERT in cells from aged individuals (**Figure 2N**). Silencing Glut1 in young endothelial cells to decrease ATP levels, also abolished the association of HSP9a with TERT (see Supplementary material online, **Figure S3E**), whilst the addition of a cell permeable ATP to cells from aged individuals re-established the TERT-Hsp9a association (see Supplementary material online, **Figure S3E**). These results indicate that reduced glycolysis and ATP levels can be directly linked with the disassociation of the active telomerase complex thus resulting in decreased telomerase activity in aged endothelial cells. To distinguish between the ATP-dependent and the telomerase dependent effects on endothelial cell proliferation, a lentiCRISPR approach was used to decrease TERT expression in native endothelial cells isolated from young individuals. Reduced TERT expression accelerated the development of replicative senescence, which became apparent after the 4^th^ cycle of passaging (**Figure 2O**). This contrasted with the situation in endothelial cells infected with the non-targeted control gRNAs that showed signs of replicative senescence only after the 10^th^ cycle of passaging. Importantly, the addition of an ATP analogue was able to increase endothelial proliferation in the multi-passaged control cells and not in TERT-deficient cells, indicating that ATP preserves proliferation only in the presence of TERT.

### Cystathionine γlyase anchors p53 in the cytoplasm to inhibit its activity

Next we addressed the mechanisms underlying the activation of p53 in aging endothelial cells and focused on the expression of cystathionine γ lyase (CSE). The reasons for this decision were (i) that 3 of the 7 most enriched metabolites (cysteine, methionine, cystathionine) detected in aged endothelial cells (see Figure 2B) could be directly linked to CSE metabolic activity, and (ii) p53 was previously identified as a CSE-associated protein.^23^

While endothleial cells from young individuals clearly expressed CSE, levels were markedly lower in aged endothelial cells (see Supplementary material online, **Figure S4A**). Similarly, there was a gradual, age-dependent decrease in endothelial CSE expression in aortic endothelial cells from 1, 6 and 12 month old wild-type mice (see Supplementary material online, **Figure S4B**) that was paralleled an increase in the acetylation of p53 on Lys120 (see Supplementary material online, **Figure S4C**). The association of p53 with CSE was initially identified by mass spectrometry,^23^ and a direct interaction between the two proteins could be demonstrated using a purified GST-CSE and recombinant p53 (**Figure 3A**). Moreover, proximity ligation and co-immunoprecipitation studies confirmed the interaction between the two proteins in the cytosol of endothelial cells from young individuals (**Figure 3B–3C**), a result that was replicated *in situ* in aorta from young but not aged individuals (**Figure 3D**).

**Figure 3.**
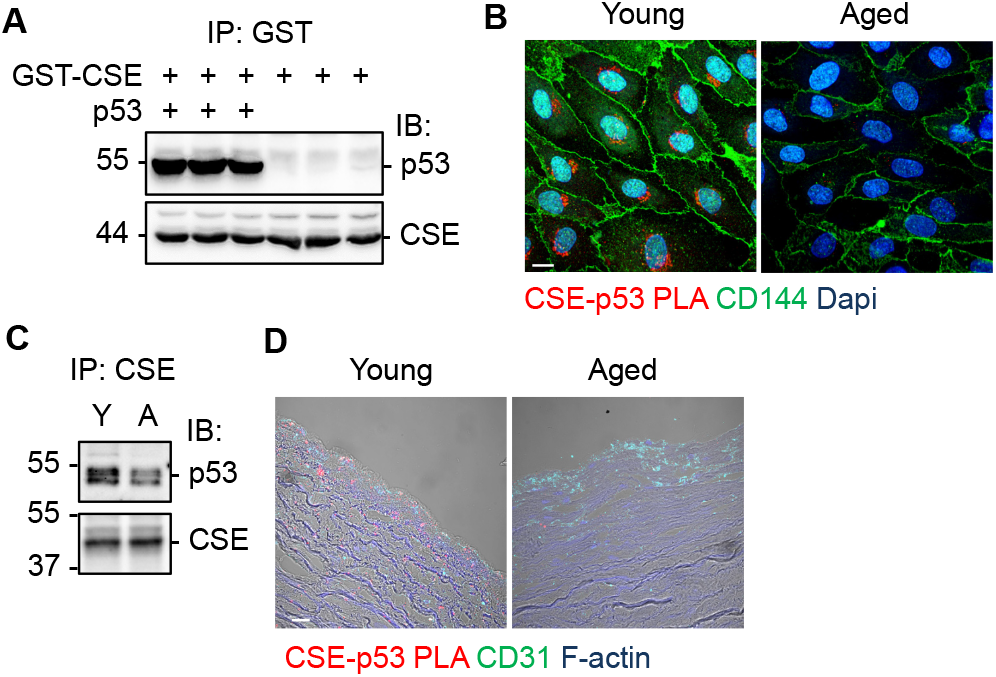
Interaction between CSE and p53 in vitro and in mesenteric artery endothelial cells isolated from young (Y) and aged (A) individuals. (**A**) Interaction of GST-CSE with p53 *in vitro*, comparable results were obtained in 2 additional experiments. (**B**) Proximity ligation assay (PLA, red) showing the interaction of CSE with p53 in CD144+ (green) young and aged endothelial cells; bar = 5 μm. Results represent 2 additional independent experiments. (**C**) Immunoblot showing p53 immunoprecipitated (IP) with CSE. Results represent 5 additional independent experiments. (**D**) Proximity ligation assay (PLA) showing the CSE-p53 interaction (red) in CD31+ (green) cells in aortae from young and aged individuals; bar = 50 μm. Results represent 3 additional independent experiments.

*In silico* docking studies predicted that the amino acid sequence Leu243, Gly244, Aal245, Val246 (LGAV) and Val95 in CSE interacted with the DNA binding site of p53 (**Figure S5A**). Indeed, the deletion of the LGAV sequence or the mutation of Val95 to lysine, attenuated the interaction of CSE with p53 (see Supplementary material online, **Figure S5B**). CSE activity generates H_2_S, which elicits its effects via the *S*-sulfhydration of target proteins.^24^ However, the interaction of p53 with CSE did not result in its *S*-sulfhydration; as determined by screening available *S*-sulfhydration datasets,^25^ and by using a p53-*S*-sulfhydration antibody array assay (see Supplementary material online, **Figure S5C**). Nor did the interaction of p53 alter CSE activity, as assessed by measuring H_2_S production (see Supplementary material online, **Figure S5D**). To determine whether or not the association with CSE could affect the nuclear translocation of p53, cells were treated with doxorubicin to induce DNA damage and p53 translocation.^26^ While doxorubicin increased nuclear levels of p53 in endothelial cells from aged individuals transduced with GFP, the overexpression of CSE resulted in the cytosolic retention of p53 (see Supplementary material online, **Figure S5E**).

The observations made in human endothelial cells were validated in murine endothelial cells i.e. CSE co-precipitated with p53 from endothelial cells from wild-type but not CSE-deficient mice (CSE^iΔEC^ mice) (see Supplementary material online, **Figure S6A-6B**). Similarly, the deletion of CSE in murine endothelial cells increased the nuclear localization of p53 (see Supplementary material online, **Figure S6C**). p53 activity is regulated by acetylation^27^ and CSE deletion enhanced the interaction between p53 with the acetyltransferase; monocytic leukaemia Zing finger (MOZ) (see Supplementary material online, **Figure S6D-6E**). The acetylation of p53 by MOZ induces senescence by promoting transactivation of the p21 promoter,^17^ and p21, in turn, mediates cell-cycle arrest in cancer cells.^20,28,29^ In murine endothelial cells lacking CSE increased expression of p53 target genes was detected (see Supplementary material online, **Figure S6F**). The binding of p53 to the Glut1 promoter was also greater in endothelial cells from CSE^IΔEC^ mice (see Supplementary material online, **Figure S6G**), resulting in reduced Glut1 expression (see Supplementary material online, **Figure S6H**), reduced uptake of 2-NBDG (Supplementary material online, **Figure S6I**), reduced flux of carbons from ^13^C-glucose to glucose-6-phosphate (see Supplementary material online, **Figure S6J**) and reduced intracellular ATP levels (see Supplementary material online, **Figure S6K**). The impact of CSE deficiency on the telomerase complex was also assessed (see Supplementary material online, **Table S3**), and the loss of CSE resulted in pronounced changes in TERT-associated proteins (see Supplementary material online, **Figure S6L**). Consistent with our findings in human endothelial cells, HSP9a was part of the telomerase complex in cells from wild-type mice but was lacking in the complex precipitated from CSE-deficient endothelial cells (see Supplementary material online, **Figure S6M**). As in the human endothelial cells, attenuated glycolysis and decreased ATP levels could be directly linked with impaired telomerase activity as the downregulation of Glut1 impaired telomerase activity in endothelial cells from wild-type mice (see Supplementary material online, **Figure S6N**). On the other hand, the attenuated telomerase activity in endothelial cells from CSE^iΔEC^ mice was increased by a cell permeable ATP analogue (see Supplementary material online, **Figure S6O**).

### Deletion of CSE results in premature endothelial cell senescence

To determine whether a change in CSE activity was a cause or consequence of vascular aging, microvascular endothelial cells were isolated from lungs of wild-type and CSE^iΛEC^ littermates. Culturing cells from CSE^iΛEC^ mice resulted in a positive SAβG signal after only 7 passages (**Figure 4A**), while approximately 25 passages were required to elicit a similar signal in endothelial cells from wild-type mice. CSE-deficient cells also proliferated more slowly (**Figure 4B**), with a higher proportion of cells accumulating in S phase (**Figure 4C**), indicating cell cycle arrest.^28^ We also observed a small but significant decrease in telomere length identified either via qPCR (**Figure 4D**) or using peptide nucleic acid fluorescent *in situ* hybridization (**Figure 4E**). Even though the expression of mRNA encoding the core telomerase subunits: TERT and telomerase RNA elements (TERC) were not altered in the absence of CSE (**Figure 4F**), telomerase activity (ddTRAP assay) was reduced (**Figure 4G**). RNA sequencing revealed the enrichment of transcripts positively regulating senescence as well as a reduction of transcripts negatively regulating senescence in cultured murine endothelial cells (passage 7) from CSE^iΔEC^ mice versus wild-type littermates (**Figure 4H**).

**Figure 4.**
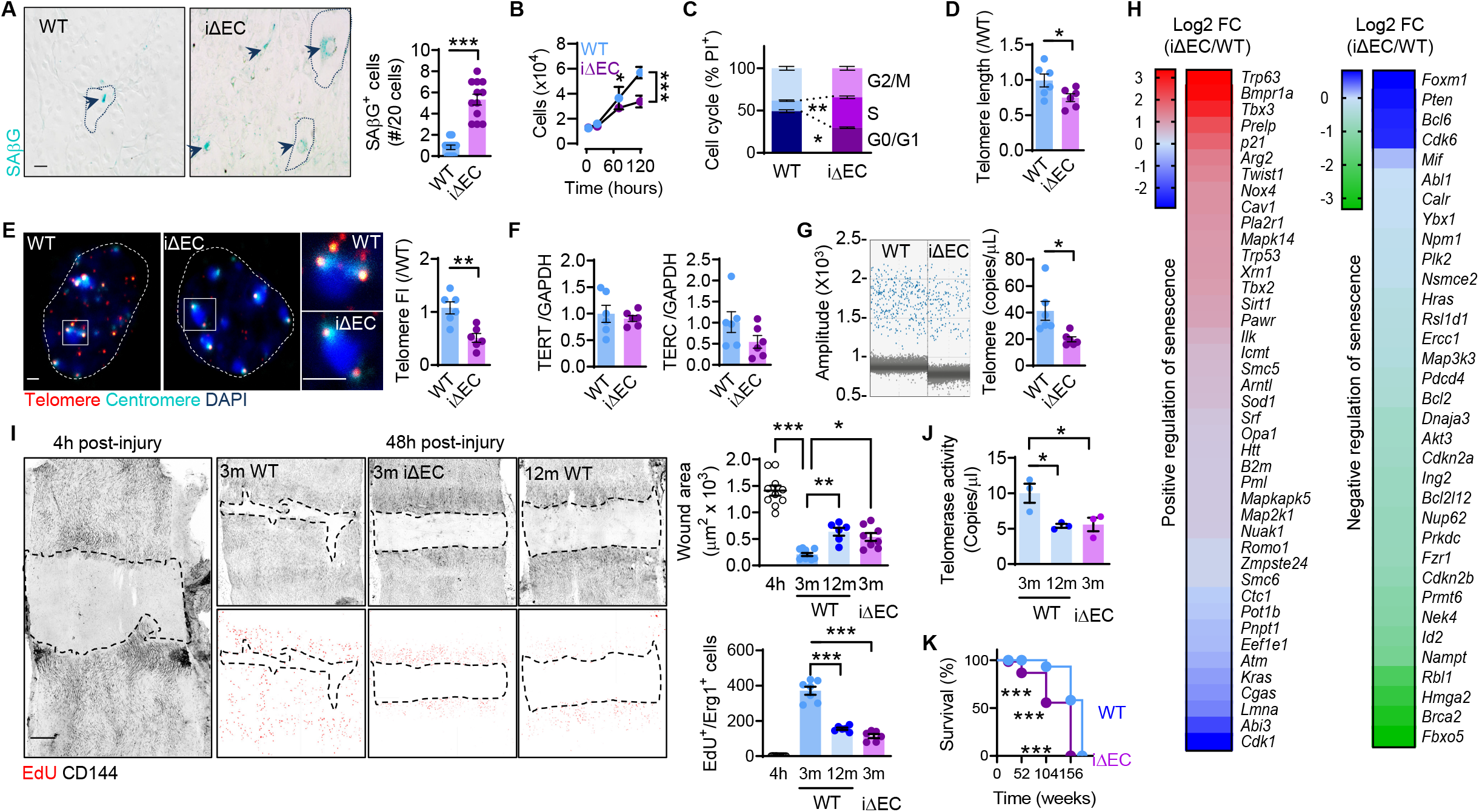
Effect of CSE deletion on endothelial cell senescence and proliferation. Pulmonary endothelial cells from wild type (WT) and CSE^iΔEC^ (iΔEC) mice were studied at passage 7; n=12/group. (**A**) Representative SAβG staining. Bar = 10 μm. (**B**) Cell proliferation over 120 hours. (**C**) Cell cycle status. (**D**) Telomere length. (**E**) Representative PNA fish imaging of telomeres (red) and centromeres (cyan), DAPI (blue). Bar=1 μm. (**F**) Relative mRNA levels of TERT and TERC normalized to GAPDH. (**G**) Representative ddTRAP image and quantification of telomerase activity. (**H**) Heatmaps showing the log2 Fold Change (FC) in positive or negative regulators of senescence, involved in cell aging (GO: 0007569). (**I**) Re-endothelization of aortae 48 hours after injury in 3 and 12 month old wild-type and 3 month old CSE^iΔEC^ mice. Proliferating cells were identified by EdU (red) incorporation. Bar = 200 μm n=6-10/group. (**J**) Telomerase activity in CD144+ murine aortic endothelial cells isolated from 3 and 12 month old wild-type mice and 3 month old CSE^iΔEC^ mice; n=3-4/group. Each sample is a pool of 3-4 animals. (**K**) Survival curve of WT and CSE^iΔEC^ mice; n=10/group. *P<0.05, **P<0.01, ***P<0.001. Student’s t-test (A, D, E, F, G), two-way ANOVA followed by Tukey’s multiple comparisons test (B, C), one-way ANOVA followed by Tukey’s multiple comparisons (I, J) or Mantel-Cox test followed by Gehan-Breslow-Wilcoxon test (K).

To assess the consequences of CSE-dependent endothelial cell senescence/proliferation *in vivo* we studied re-endothelization 48 hours after injury of the abdominal aorta by assessing wound area coverage and EdU positive cell numbers. Consistent with a previous report,^29^ endothelial cell regeneration after injury was rapid and approximately 80% of the wounded area was re-endothelialized after 48 hours in 3 month old wild-type mice. In 3 month old CSE^iΔEC^ mice repopulation of the damaged area was much slower and was comparable to that recorded in 12 month old wild-type mice (**Figure 4I**). Consistent with this observation telomerase activity was significantly reduced in cells from CSE^iΛEC^ (**Figure 4J**), indicating that CSE deletion resulted in accelerated vascular aging. Indeed, the life span of CSE^iΛEC^ mice was shorter than that of their CSE-expressing littermates (**Figure 4K**).

*In vitro* rescue experiments in which murine endothelial cells lacking CSE (passage 7) or wild type endothelial cells (passage 25) were adenovirally transduced to overexpress CSE resulted in the loss of SAβG positive staining (see Supplementary material online, **Figure S7A**), increased telomerase activity (Supplementary material online, **Figure S7B**) resulted in longer telomeres (see Supplementary material online, **Figure S7C**) and stimulated cell cycle progression (see Supplementary material online, **Figure S7D**). The adenoviral mediated re-introduction of CSE into CSE-deficient endothelial cells, decreased the binding of p53 to the Glut1 promoter (**Figure 5A**), and increased Glut 1 expression on the cell surface (**Figure 5B**). This approach also activated glycolysis (**Figure 5C**), enhanced glycolytic capacity (**Figure 5D**), increased ATP levels (**Figure 5E**) and restored the interaction of Hsp9a with telomerase (**Figure 5F**). Thus, the ablation of CSE in murine endothelial cells resulted in the rapid development of a replicative senescence phenotype that was similar to that detected in native endothelial cells from aged individuals.

**Figure 5.**
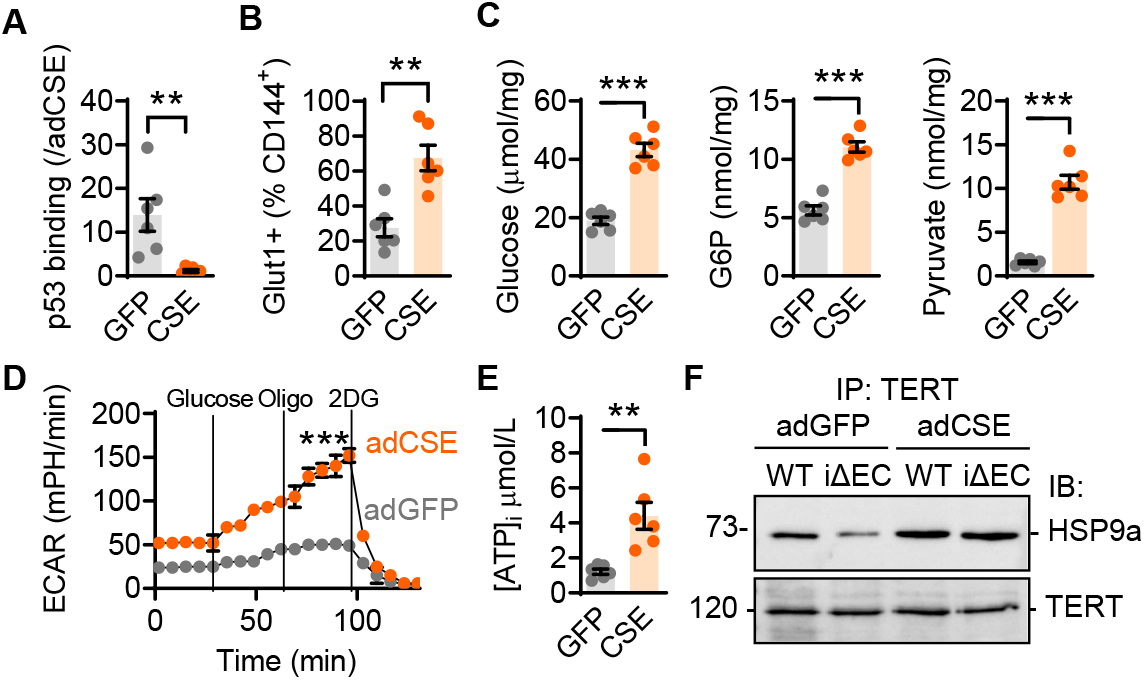
Consequences of CSE overexpression on glycolysis and the telomerase complex. Endothelial cells from CSE^iΔEC^ mice were transduced with adenovirsuses encoding GFP or CSE from passage 2 up to passage 7. Cells were studied after passage 7; n=6/group. (**A**) p53 binding to the Glut1 promoter identified by ChiP-qPCR. (**B**) Glut1 positive cell populations. (**C**) Glucose, glucose-6 phosphate (G6P) and pyruvate levels. (**D**) Extracellular acidification rate (ECAR) as an index of glycolysis. (**E**) Intracellular (i) ATP levels. (**F**) Association of Hsp9a with TERT. Comparable observations were made in 5 additional experiments. **P<0.01, ***P<0.001. Student’s t-test (A, B, C, E) and two-way ANOVA followed Tukey’s multiple comparisons test (D).

**Figure 6.**
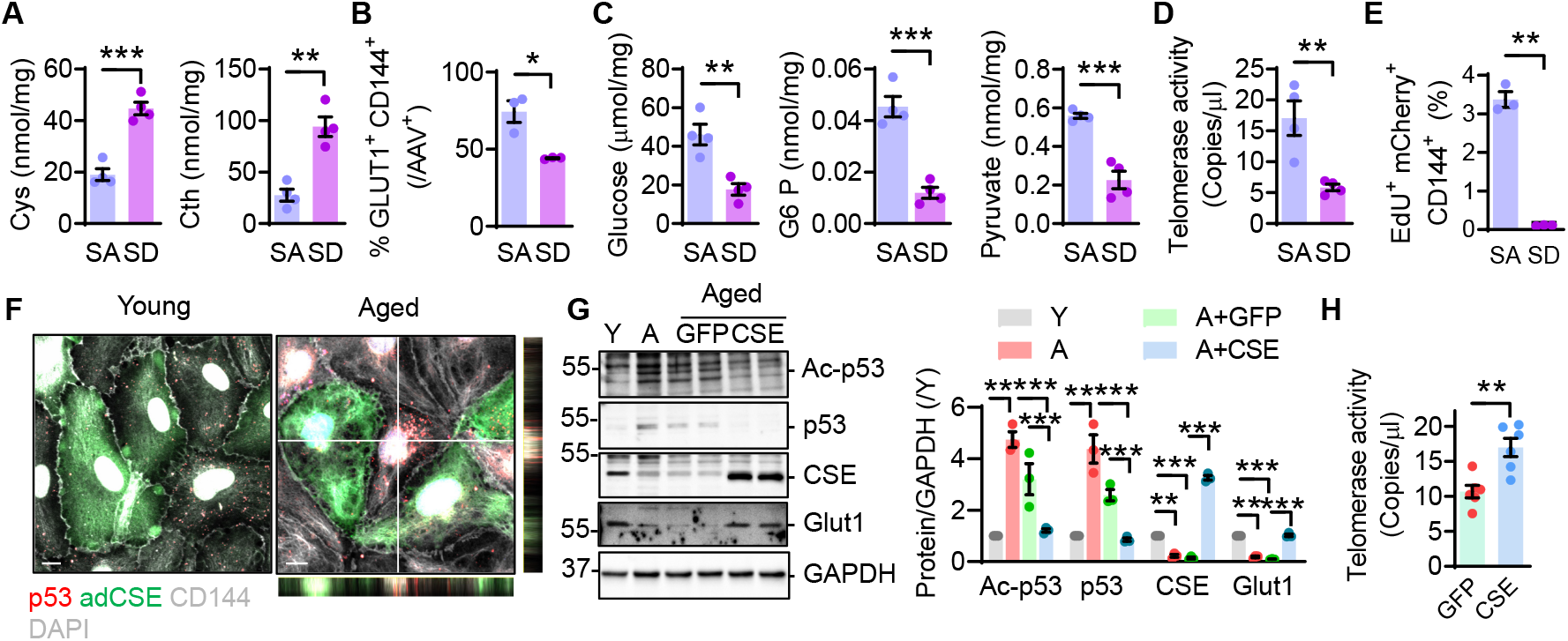
CSE reexpression rescues endothelial cell senescense. (**A-E**) 3 month old CSE^iΔEC^ mice were injected with AAV9 particles encoding constitutively active CSE-S377A-mCherry (SA) or inactive CSE-S377D-mCherry (SD). CD144+ mCherry+ cells were isolated after 9 months (FACS sorting). (A) Cysteine (Cys) and cystathionine (Cth) levels, (B) Glut 1 surface expression. (C) Glucose, glucose-6 phosphate (G6P) and pyruvate levels. (D) Telomerase activity. (E) CD144+ EdU+ mCherry+ cells from mice subjected to aortic regeneration; n=3-4/group with each sample a pool of 3 animals. (**F-H**) Human native endothelial cells from young or aged subjects were transduced using a CSE-coding adenovirus (10 MOI, 6 days). (F) Images showing p53 (red), adCSE (green) and DAPI, CD144 (grey), comparable results were observed in 2 additional cell isolates. Bar = 5 μm. (G) Immunoblots for acetylated p53 (at K120), p53, p21 and Glut1; n=3/group. (H) Telomerase activity in aged endothelial cells lacking (GFP) or expressing adCSE (CSE); n=3-6/group. *P<0.05, **P<0.01, ***p<0.001 Student’s t-test (A-E, H), one way ANOVA and Tukey’s multiple comparisons test (G).

### CSE expression as a novel approach to reduce p53 activity, preserve telomerase activity and delay endothelial cell senescence

Next we tested the hypothesis that the re-expression of CSE in senescent endothelial cells could rescue endothelial cell proliferation and vascular repair, using endothelial cell-targeted adeno-associated 9 (AAV9) viruses to express either a constitutively active CSE mutant i.e. CSE- S377A-mCherry, or a catalytically inactive mutant i.e. CSE-S377D-mCherry. Here it is important to note that the inactive S377D mutant bound less p53 (~ 50% less) than the constitutively active mutant (see Supplementary material online, **Figure S8A**). Although the incorporation of the CSE mutants was very low, it was possible to demonstrate that the metabolic profile of mCherry+ aortic endothelial cells was partially normalized by S377A-CSE (Supplementary material online, **Figure S8B**). Also, compared with CSE-S377D, CSE-S377A-expressing cells consumed more cysteine and cystathionine i.e had a higher CSE activity (**Figure 6A**), expressed more Glut1 on the cell surface (**Figure 6B**), had higher intracellular glucose levels (**Figure 6C**, see Supplementary material online, **Figure S8C**), and demonstrated increased telomerase activity (**Figure 6D**). The rescue also improved proliferation as endothelial cells from mice that expressed the S377A-CSE mutant prior to aortic injury were EdU positive 48 hours after the intervention. No EdU positive signal was detected in endothelial cells expressing S377D-CSE (**Figure 6E**). Unfortunately, the limited efficiency of viral transduction meant that it was not possible to draw conclusions about alterations in endothelial regenerative capacity. Therefore, we determined the consequences of reintroducing CSE into endothelial cells from young and aged individuals which were reintroduced into culture for 6 days (without passaging). The CSE construct used also contained GFP to facilitate the detection of transduced cells. In cells from young individuals p53 expression was detectable and mostly cytosolic and the overexpression of CSE decreased p53 levels (**Figure 6F**); probably by inactivating HuR.^23^ In the aged endothelial cells p53 levels were higher and the protein was detected in the cytosol as well as the nucleus. In these cells, CSE overexpression decreased the nuclear localization of p53. Additionaly, p53 acetylation and the expression of p21 were lower in CSE-transduced, aged endothelial cells, while Glut1 expression was increased (**Figure 6G**). A significant increase in telomerase activity was also detected in the aged human endothelial cells following the reintroduction of CSE (**Figure 6H**).

## Discussion

In our study we used a unique collection of native human endothelial cells from young and old individuals to demonstrate that endothelial cell senescence is linked to the loss of a previously described vasoprotective enzyme i.e. CSE, which physically interacts with p53 to retain it in the cytosol and prevent its activation. Indeed, the endothelial cell-specific deletion of CSE in mice resulted in premature cell senescence, concomitant with the nuclear accumulation of p53, its acetylation and the initiation of a pro-senescence gene program. There were also tight links to metabolic reprograming as the loss of CSE and activation of p53 reduced Glut 1 expression, glucose transport, glycolysis and ATP levels to such an extent that the ATP-dependent chaperone Hsp9a was unable to bind to TERT and form the active telomerase complex. These observations are relevant inasmuch as according to the *“metabolic telomere attrition”* hypothesis, altered metabolism results in swift reorganization of the telomere complex to result in telomere attrition.^30^

Senescent cells can impact on homeostasis by limiting the regenerative capacity of a tissue and using a previously described model of murine aortic injury,^29^ we were able to show that aging impairs endothelial cell regeneration in mice. Moreover, even though murine telomeres are longer than human telomeres,^31^ reduced endothelial cell proliferation was linked to a reduction in telomere length as well as to impaired glycolysis. Importantly, the attenuated recovery observed in 12 month old wild-type mice was phenocopied in 3 month old endothelial cell-specific CSE knockout mice and partially rescued by the reintroduction of a CSE-S377A mutant that bound p53 more effectively than an inactive CSE-S377D mutant. Although the transduction efficiency was not sufficient to promote accelerated re-endothelization, endothelial cells expressing the active CSE mutant exhibited increased proliferation, preserved glycolytic flux and telomerase activity, showing resistance to natural aging.

Numerous studies have assessed the impact of p53 in senescence and longevity.^32–34^ In non- vascular cells the p53 protein mainly localizes to the cytoplasm and its rapid ubiquitination and turnover maintains a low level of expression.^35^ In response to prolonged cellular stress, the p53 protein is stabilized as a result of increased expression of the main p53 E3 ubiquitin ligase; MDM2.^35,36^ Although, the mechanisms regulating p53 expression and translocation in aging endothelial cells have not been studied in detail, we observed a clear age-associated increase nuclear p53 levels as well as p53-dependent transcriptional activity that, in native endothelial cells from aged individuals, coincided increased expression of MDM2. Other mechanisms have been proposed to regulate the subcellular localization of p53 including its association with cytosolic proteins, one example being its ability of interact with Foxo4 in fibroblasts.^37^ The results of our study have identified CSE as an additional p53 trap in endothelial cells and identified 3 sites within the CSE proteins that determine the interaction between the two proteins i.e. the LGAV sequence (Leu243-Val246), Val95 and Ser377. The latter serine residue was previously identified as a critical regulator of CSE activity, and its phosphorylation in response to endothelial cell exposure to inflammatory cytokines results in rapid enzyme inactivation during atherogenesis.^23^ The finding that the phosphorylated enzyme also binds p53 less effectively implies that vascular inflammation and endothelial cells activation also accelerate endothelial cell senescence.

Ours is not the first report to link altered cystathionine and cysteine metabolism with cell survival or longevity. Indeed, H_2_S can *S*-sulfhydrate proteins and higher protein *S*-sulfhydration levels induced by pharmacological or dietary interventions, lead to better resistance to oxidative stress and longer life.^38–41^ Also, H_2_S generation has been linked with the beneficial effects of calorie restriction (sulfur amino acid restriction) on lifespan.^42,43^ However, p53 was not directly targeted by H_2_S and the protein was not *S*-sulfhydrated in human endothelial cells in either this study or our previous investigation.^25^ Moreover, the interaction of the p53 with CSE was independent of CSE enzymatic activity. While there is no doubt that CSE-derived H_2_S largely determines the *S*-sulfhydration of endothelial cell proteins,^25^ many of which are involved in the regulation of endothelial function and the cellular redox environment, it is very unlikely that the exogenous application of H_2_S would be able to reverse the aging process. Indeed,the nucleophilic cysteine residues normally targeted by H_2_S tend to be irreversibly *S*-sulfenylated in aged cells; a modification that replaces the protective *S*-sulfhydration.^39^ This implies that while H_2_S may protect against the occurrence of protein hyperoxidation, as is the case for peroxiredoxin 6,^44^ H_2_S would not be able to re-sulfhydrate already sulfenylated targets to elicit cellular rejuvenation. In contrast, approaches to increase CSE expression would result in the physical retention of p53 in the cytosol to dampen its activation and delay the p53-dependent aging process. The potential effectiveness of such a stragegy was demonstrated using an AAV-based approach to reintroduce CSE into either aged or CSE-deficient endothelial cells which resulted in the capture cytosolic p53, a decrease in its nuclear translocation and activation, and the preservation of glycolytic flux to maintain a high energetic endothelial state with reduced senescence. Thus, strategies to prevent the age-dependent decline in CSE expression or to reintroduce CSE into endothelial cells would be an attractive approach to prevent the metabolic impairment of telomerase activity and delay endothelial cell senescence and the onset of cardiovascular disease.

## Data Availability

The authors declare that the data supporting the findings of this study are available within the paper and its Supplementary Material.

## Acknowledgements

The authors are indebted to Isabel Winter, Cindy Fabiene Hoeper, Katharina Herbig and Oliver Haun for expert technical assistance.

## Author Contributions

S.-I.B., J.H. and I.F. designed the project. S.-I.B. and I.F. wrote the paper. S.-I.B., J.H., M.S.L., G.D., M.S., J.W., M.-K.D., S.Z., J.O., A.W. generated data. S.-I.B., J.H., A.W., and M.L., analyzed the data. S.G., M.L. and D.J. analyzed the sequencing data. I.W. performed the proteomic analysis. S.H. and O.-J.M. generated the AAVs, A.P. and F.S. collected and analyzed the human data. All authors read the manuscript and provided feedback before submission.

---

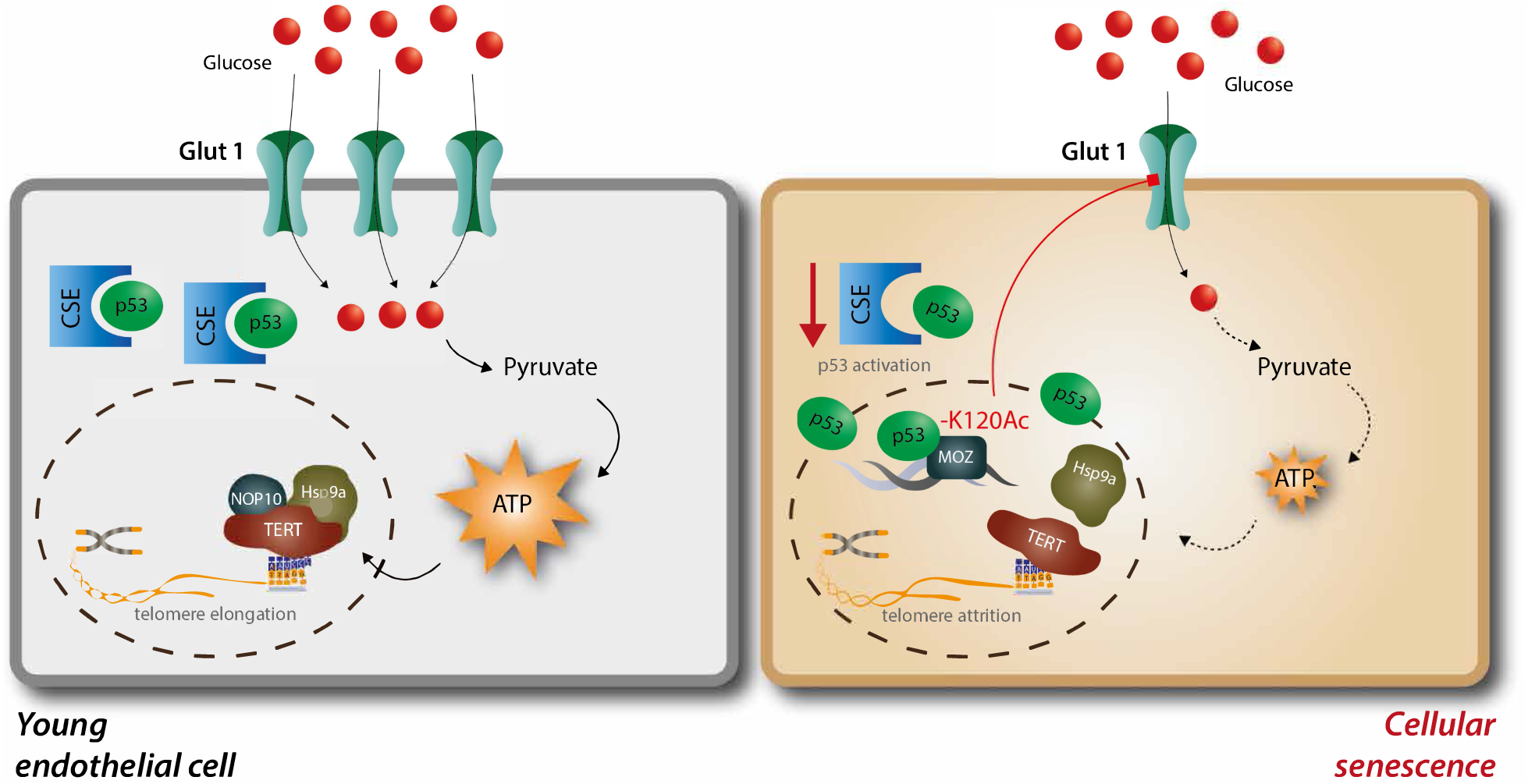

## References

1. Ungvari Z, Tarantini S, Donato AJ, Galvan V, Csiszar A. Mechanisms of vascular aging. Circ Res 2018;123:849–867.

2. Regnault V, Challande P, Pinet F, Li Z, Lacolley P. Cell senescence: basic mechanisms and the need for computational networks in vascular ageing. Cardiovasc Res 2021;117:1841–1858.

3. Nagai Y, Metter EJ, Earley CJ, Kemper MK, Becker LC, Lakatta EG, Fleg JL. Increased carotid artery intimal-medial thickness in asymptomatic older subjects with exercise-induced myocardial ischemia. Circulation 1998;98:1504–1509.

4. Franklin SS, Khan SA, Wong ND, Larson MG, Levy D. Is pulse pressure useful in predicting risk for coronary heart disease? The Framingham heart study. Circulation 1999;100:354–360.

5. Campisi J. Aging, cellular senescence, and cancer. Ann Rev Physiol 2013;75:685–705.

6. Beauséjour CM, Krtolica A, Galimi F, Narita M, Lowe SW, Yaswen P, Campisi J. Reversal of human cellular senescence: roles of the p53 and p16 pathways. EMBO J 2003;22:4212–4222.

7. Baker DJ, Childs BG, Durik M, Wijers ME, Sieben CJ, Zhong J, Saltness RA, Jeganathan KB, Verzosa GC, Pezeshki A, Khazaie K, Miller JD, van Deursen JM. Naturally occurring p16(Ink4a)-positive cells shorten healthy lifespan. Nature 2016;530:184–189.

8. Xu M, Pirtskhalava T, Farr JN, Weigand BM, Palmer AK, Weivoda MM, Inman CL, Ogrodnik MB, Hachfeld CM, Fraser DG, Onken JL, Johnson KO, Verzosa GC, Langhi LGP, Weigl M, Giorgadze N, LeBrasseur NK, Miller JD, Jurk D, Singh RJ, Allison DB, Ejima K, Hubbard GB, Ikeno Y, Cubro H, Garovic VD, Hou X, Weroha SJ, Robbins PD, Niedernhofer LJ, Khosla S, Tchkonia T, Kirkland JL. Senolytics improve physical function and increase lifespan in old age. Nat Med 2018;24:1246–1256.

9. Suda M, Shimizu I, Katsuumi G, Yoshida Y, Hayashi Y, Ikegami R, Matsumoto N, Yoshida Y, Mikawa R, Katayama A, Wada J, Seki M, Suzuki Y, Iwama A, Nakagami H, Nagasawa A, Morishita R, Sugimoto M, Okuda S, Tsuchida M, Ozaki K, Nakanishi-Matsui M, Minamino T. Senolytic vaccination improves normal and pathological age-related phenotypes and increases lifespan in progeroid mice. Nat Aging 2021;1:1117–1126.

10. Asai K, Kudej RK, Shen YT, Yang GP, Takagi G, Kudej AB, Geng YJ, Sato N, Nazareno JB, Vatner DE, Natividad F, Bishop SP, Vatner SF. Peripheral vascular endothelial dysfunction and apoptosis in old monkeys. Arterioscler Thromb Vasc Biol 2000;20:1493–1499.

11. Haudenschild CC, Prescott MF, Chobanian AV. Aortic endothelial and subendothelial cells in experimental hypertension and aging. Hypertension 1981;3:I148–53.

12. Li Z, Froehlich J, Galis ZS, Lakatta EG. Increased expression of matrix metalloproteinase-2 in the thickened intima of aged rats. Hypertension 1999;33:116–123.

13. Marchand A, Atassi F, Gaaya A, Leprince P, Le Feuvre C, Soubrier F, Lompré A-M, Nadaud S. The Wnt/beta-catenin pathway is activated during advanced arterial aging in humans. Aging Cell 2011;10:220–232.

14. Coufal J, Jagelská EB, Liao JCC, Brázda V. Preferential binding of p53 tumor suppressor to p21 promoter sites that contain inverted repeats capable of forming cruciform structure. Biochem Biophys Res Com 2013;441:83–88.

15. Chen J, Huang X, Halicka D, Brodsky S, Avram A, Eskander J, Bloomgarden NA, Darzynkiewicz Z, Goligorsky MS. Contribution of p16INK4a and p21CIP1 pathways to induction of premature senescence of human endothelial cells: permissive role of p53. Am J Physiol Heart Circ Physiol 2006;290:H1575–H1586.

16. Warboys CM, Luca A de, Amini N, Le Luong, Duckles H, Hsiao S, White A, Biswas S, Khamis R, Chong CK, Cheung W-M, Sherwin SJ, Bennett MR, Gil J, Mason JC, Haskard DO, Evans PC. Disturbed flow promotes endothelial senescence via a p53-dependent pathway. Arterioscler Thromb Vasc Biol 2014;34:985–995.

17. Reed SM, Quelle DE. p53 Acetylation: regulation and consequences. Cancers (Basel) 2014;7:30–69.

18. Kruiswijk F, Labuschagne CF, Vousden KH. p53 in survival, death and metabolic health: a lifeguard with a licence to kill. Nat Rev 2015;16:393–405.

19. Puzio-Kuter AM. The role of p53 in metabolic regulation. Genes Cancer 2011;2:385–391.

20. Schwartzenberg-Bar-Yoseph F, Armoni M, Karnieli E. The tumor suppressor p53 down-regulates glucose transporters GLUT1 and GLUT4 gene expression. Cancer Res 2004;64:2627–2633.

21. Depcrynski AN, Sachs PC, Elmore LW, Holt SE. Regulation of telomerase through transcriptional and posttranslational mechanisms. In: Hiyama K, editor. Telomeres and Telomerase in Cancer. Totowa, NJ: Humana Press, 2009:47–85.

22. Forsythe HL, Jarvis JL, Turner JW, Elmore LW, Holt SE. Stable association of hsp90 and p23, but Not hsp70, with active human telomerase. J Biol Chem 2001;276:15571–15574.

23. Bibli S-I, Hu J, Sigala F, Wittig I, Heidler J, Zukunft S, Tsilimigras DI, Randriamboavonjy V, Wittig J, Kojonazarov B, Schürmann C, Siragusa M, Siuda D, Luck B, Abdel Malik R, Filis KA, Zografos G, Chen C, Wang DW, Pfeilschifter J, Brandes RP, Szabo C, Papapetropoulos A, Fleming I. Cystathionine γ lyase sulfhydrates the RNA binding protein human antigen R to preserve endothelial cell function and delay atherogenesis. Circulation 2019;139:101–114.

24. Bibli S-I, Fleming I. Oxidative post-translational modifications: A focus on cysteine S-sulfhydration and the regulation of endothelial fitness. Antioxid Redox Signal 2021;35:1494–1514.

25. Bibli S-I, Hu J, Looso M, Weigert A, Ratiu C, Wittig J, Drekolia MK, Tombor L, Randriamboavonjy V, Leisegang MS, Goymann P, Delgado Lagos F, Fisslthaler B, Zukunft S, Kyselova A, Justo AFO, Heidler J, Tsilimigras D, Brandes RP, Dimmeler S, Papapetropoulos A, Knapp S, Offermanns S, Wittig I, Nishimura SL, Sigala F, Fleming I. Mapping the endothelial cell S-sulfhydrome highlights the crucial role of integrin sulfhydration in vascular function. Circulation 2021;143:935–948.

26. Saha T, Guha D, Manna A, Panda AK, Bhat J, Chatterjee S, Sa G. G-actin guides p53 nuclear transport: potential contribution of monomeric actin in altered localization of mutant p53. Sci Rep 2016;6:32626.

27. Barlev NA, Liu L, Chehab NH, Mansfield K, Harris KG, Halazonetis TD, Berger SL. Acetylation of p53 activates transcription through recruitment of coactivators/histone acetyltransferases. Mol Cell 2001;8:1243–1254.

28. Zilfou JT, Lowe SW. Tumor suppressive functions of p53. Cold Spring Harb Perspect Biol 2009;1:a001883.

29. McDonald AI, Shirali AS, Aragón R, Ma F, Hernandez G, Vaughn DA, Mack JJ, Lim TY, Sunshine H, Zhao P, Kalinichenko V, Hai T, Pelegrini M, Ardehali R, Iruela-Arispe ML. Endothelial regeneration of large vessels is a biphasic process driven by local cells with distinct proliferative capacities. Cell Stem Cell 2018;23:210–225.e6.

30. Casagrande S, Hau M. Telomere attrition: metabolic regulation and signalling function? Biol Lett 2019;15:20180885.

31. Vera E, Bernardes de Jesus B, Foronda M, Flores JM, Blasco MA. The rate of increase of short telomeres predicts longevity in mammals. Cell Rep 2012;2:732–737.

32. Fitzgerald AL, Osman AA, Xie T-X, Patel A, Skinner H, Sandulache V, Myers JN. Reactive oxygen species and p21Waf1/Cip1 are both essential for p53-mediated senescence of head and neck cancer cells. Cell Death Dis 2015;6:e1678.

33. Kim J, Nakasaki M, Todorova D, Lake B, Yuan C-Y, Jamora C, Xu Y. p53 Induces skin aging by depleting Blimp1+ sebaceous gland cells. Cell Death Dis 2014;5:e1141.

34. Fatt MP, Cancino GI, Miller FD, Kaplan DR. p63 and p73 coordinate p53 function to determine the balance between survival, cell death, and senescence in adult neural precursor cells. Cell Death Differ 2014;21:1546–1559.

35. Marchenko ND, Hanel W, Li D, Becker K, Reich N, Moll UM. Stress-mediated nuclear stabilization of p53 is regulated by ubiquitination and importin-alpha3 binding. Cell Death Differ 2010;17:255–267.

36. Marine J-C. p53 stabilization: the importance of nuclear import. Cell Death Differ 2010;17:191–192.

37. Baar MP, Brandt RMC, Putavet DA, Klein JDD, Derks KWJ, Bourgeois BRM, Stryeck S, Rijksen Y, van Willigenburg H, Feijtel DA, van der Pluijm I, Essers J, van Cappellen WA, van IJcken WFJ, Houtsmuller AB, Pothof J, Bruin RWF de, Madl T, Hoeijmakers JHJ, Campisi J, Keizer PLJ de. Targeted apoptosis of senescent cells restores tissue homeostasis in response to chemotoxicity and aging. Cell 2017;169:132–147.e16.

38. Das A, Huang GX, Bonkowski MS, Longchamp A, Li C, Schultz MB, Kim L-J, Osborne B, Joshi S, Lu Y, Treviño-Villarreal JH, Kang M-J, Hung T-T, Lee B, Williams EO, Igarashi M, Mitchell JR, Wu LE, Turner N, Arany Z, Guarente L, Sinclair DA. Impairment of an endothelial NAD+-H2S signaling network is a reversible cause of vascular aging. Cell 2018;173:74–89.e20.

39. Zivanovic J, Kouroussis E, Kohl JB, Adhikari B, Bursac B, Schott-Roux S, Petrovic D, Miljkovic JL, Thomas-Lopez D, Jung Y, Miler M, Mitchell S, Milosevic V, Gomes JE, Benhar M, Gonzalez-Zorn B, Ivanovic-Burmazovic I, Torregrossa R, Mitchell JR, Whiteman M, Schwarz G, Snyder SH, Paul BD, Carroll KS, Filipovic MR. Selective persulfide detection reveals evolutionarily conserved antiaging effects of S-sulfhydration. Cell Metab 2019;30:1152–1170.e13.

40. Statzer C, Meng J, Venz R, Bland M, Robida-Stubbs S, Patel K, Petrovic D, Emsley R, Liu P, Morantte I, Haynes C, Mair WB, Longchamp A, Filipovic MR, Blackwell TK, Ewald CY. ATF-4 and hydrogen sulfide signalling mediate longevity in response to inhibition of translation or mTORC1. Nat Commun 2022;13:967.

41. Wilkie SE, Borland G, Carter RN, Morton NM, Selman C. Hydrogen sulfide in ageing, longevity and disease. Biochem J 2021;478:3485–3504.

42. Hine C, Mitchell JR. Calorie restriction and methionine restriction in control of endogenous hydrogen sulfide production by the transsulfuration pathway. Exp Gerontol 2015;68:26–32.

43. Hine C, Harputlugil E, Zhang Y, Ruckenstuhl C, Lee BC, Brace L, Longchamp A, Treviño-Villarreal JH, Mejia P, Ozaki CK, Wang R, Gladyshev VN, Madeo F, Mair WB, Mitchell JR. Endogenous hydrogen sulfide production is essential for dietary restriction benefits. Cell 2015;160:132–144.

44. Bibli S-I, Hu J, Leisegang MS, Wittig J, Zukunft S, Kapasakalidi A, Fisslthaler B, Tsilimigras D, Zografos G, Filis K, Brandes RP, Papapetropoulos A, Sigala F, Fleming I. Shear stress regulates cystathionine γ lyase expression to preserve endothelial redox balance and reduce membrane lipid peroxidation. Redox Biol 2020;28:101379.

