## Supplementary Material and Methods for "A novel role for cystathionine γ lyase in the control of p53: impact on endothelial senescence and metabolic reprograming"

### METHODS

**Data Availability:** The authors declare that the data supporting the findings of this study are available within the paper and its Supplementary Information.

**Materials**

Cell culture media were from Gibco (Invitrogen; Darmstadt, Germany), OCT Tissue Tek was from Sakura (Staufen, Germany) and the protease inhibitor cocktail was from Roche (Mannheim, Germany). Sulfane Sulfur Probe 4 (SSP4) was from Dojindo (GERBU Biotechnik GmbH, Heidelberg, Germany). Fibronectin was from Corning (Kaiserslautern, Germany), DyLight 488 NHS ester was from Thermo Scientific (Karlsruhe, Germany). Copper II, TBTA was from Lumiprobe (Hannover, Germany) and Daz2 was from Cayman Chemical (Michigan, USA). All other chemicals (unless otherwise specified), were from Millipore, Sigma (Darmstadt, Germany).

**Antibodies**

All the secondary antibodies for Western blotting were from Calbiochem (Bad Soden am Taunus, Germany), and Alexa-Fluor antibodies for immunohistochemistry were from Invitrogen (Karlsruhe, Germany). All antibodies were validated either using appropriate plasmids or positive/negative controls or by the manufacturers. Antibodies used for immunohistochemistry and FACS were validated by using primary antibodies alone, secondary antibodies alone or with the blocking peptides/isotype controls. The following antibodies were used:

| Antibody | Catalogue # | Company | Dilution |
| --- | --- | --- | --- |
| **Western blotting/ immunoprecipitation** | | | |
| eNOS | 610297 | BD (Heidelberg, Germany) | 1:1000 |
| β-actin | MAK6019 | Linaris (Dossenheim, Germany) | 1:5000 |
| CSE | 12217-1-AP | Proteintech ( Rosemont, USA) | 1:500 |
| CBS | H00000875-M01 | Abnova (Heidelberg, Germany) | 1:1000 |
| 3MST | HPA001240 | Atlas (Bromma, Sweden) | 1:2000 |
| P53 | 90001-MM06 | SinoBiological (Eschborn, Germany) | 1:1000 |
| P53 (for human cells) | 9282 | Cell Signaling Technologies (Danvers, MA, USA) | 1:1000 |
| Histone 3 | 39763 | Active Motif (Waterloo, Belgium) | 1:1000 |
| GAPDH | MAB374 | Millipore (Darmstadt, Germany) | 1:2000 |
| Acetylated p53 K120 | Ab78316 | Abcam (Berlin, Germany) | 1:1000 |
| Tip60 | 12058 | Cell Signaling Technologies (Danvers, MA, USA) | 1:1000 |
| MOF (MYST1) | 46862 | Cell Signaling Technologies (Danvers, MA, USA) | 1:1000 |
| MOZ (MYST3) | 78462 | Cell Signaling Technologies (Danvers, MA, USA) | 1:1000 |
| P21 | 2947 | Cell Signaling Technologies (Danvers, MA, USA) | 1:1000 |
| Glut 1 | 12939 | Cell Signaling Technologies (Danvers, MA, USA) | 1:1000 |
| HK1 | 2024 | Cell Signaling Technologies (Danvers, MA, USA) | 1:1000 |
| HK2 | 2867 | Cell Signaling Technologies (Danvers, MA, USA) | 1:1000 |
| PKM2 | 4053 | Cell Signaling Technologies (Danvers, MA, USA) | 1:1000 |
| LDH | 3582 | Cell Signaling Technologies (Danvers, MA, USA) | 1:1000 |
| TERT | 582005 | Merck (Darmstadt, Germany) | 1:2000 |
| Hsp9a (GRP 75) | sc-133137 | Santa Cruz Biotechnology (Heidelberg, Germany) | 1:1000 |
| **ChiP antibody** | | | |
| anti-p53 | 645703 | Biolegend (Greenwood Place, London) | 1 µg/sample |
| **Immunohistochemistry/ PLA** | | | |
| CSE | 12217-1-AP | Proteintech (Rosemont, USA) | 1:400 |
| CD144 anti-mouse APC | 138012 | Biolegend (Greenwood Place, London) | 1:100 |
| P53 | 90001-MM06 | SinoBiological (Eschborn, Germany) | 1:500 |
| MOZ (MYST3) | 78462 | Cell Signaling Technologies (Danvers, MA, USA) | 1:500 |
| TERT | 582005 | Millipore (Darmstadt, Germany) | 1:500 |
| Hsp9a (GRP 75) | sc-133137 | Santa Cruz Biotechnology (Heidelberg, Germany) | 1:500 |
| CD31 | 550274 | BD Biosciences (Heidelberg, Germany) | 1:100 |
| Alexa Fluor 546 Phalloidin | A22283 | ThermoFischer (Dreieich, Germany) | 1:5000 |
| **FACS** | | | |
| Propidium iodide | 556463 | BD Biosciences (Heidelberg, Germany) | 1:100 |
| CD144 anti-mouse APC | 138012 | Biolegend (Greenwood Place, London) | 1:100 |
| CD144 anti- human APC | 348507 | Biolegend (Greenwood Place, London) | 1:50 |

**Endothelial cell isolation and culture**

*Human endothelial cells from mesenteric arteries*. Healthy mesenteric arteries from humans of 20±2.3 and 80±3.4 years of age (**Table S4**), were isolated and stored in EGM-2 media (Lonza, Cologne, Germany) supplemented with 5% orthologous human serum for 30-45 minutes. Thereafter, arteries were incubated with dispase (5 Units/mL) for 20 minutes in Hams medium (Lonza, Cologne, Germany) without serum. Subsequently, cells were pelleted and labelled with anti-CD144 antibody. Cells were then subjected to FACS sorting with a BD FACSAria III (BD Biosciences, Germany). CD144 positive cells were collected in EGM-2 media containing orthologous human serum (5%) and incubated for 15 minutes in a humidified chamber (21% O_2_, 5% CO_2_) before being harvested by centrifugation (300g, 4 minutes) and either snap frozen in liquid N_2_ until evaluation, or cryopreserved in 10% DMSO in EGM-2 media for functional analyses. The isolation procedure was approved by the Scientific and Ethic Committee of Hipokrateion University Hospital (extension to SC55/22-2-2018) and the Goethe University. In cases that native cells were cultured, endothelial cells were seeded in a mixed matrix of laminin, gelatin and fibronectin (each 2.5 ug/ml) coated dishes and cultured with ECGM-2 media (Lonza, Cologne, Germany) for the appropriate time.

*Murine aortic endothelial cells.* Murine aortae were dissected from 1, 6 and 12 month old mice and maintained in DMEM/F12 (Lonza, Cologne, Germany) supplemented with orthologous mouse serum (5%) for 15-30 minutes. Subsequently, aortas were incubated with dispase (5 Units/mL) for 20 minutes in Hams medium without serum, and endothelial cells were recovered by centrifugation (300g, 4 minutes) and labelled with anti-CD144 antibodies. Cells were subjected to FACS sorting with a BD FACSAria III (BD Biosciences, Germany). CD144 positive cells were collected in DMEM/F12 media containing orthologous mouse serum (5%) and allowed to recover for 15 minutes in a humidified chamber (21% O_2_, 5% CO_2_), before being centrifuged (300g, 4 minutes) and snap frozen in liquid N_2_ until evaluation.

*Murine lung endothelial cells*. Cells were isolated from either wild-type or CSE^iΔEC^ mice, cultured as described,^1^ and used between passages 4 and 7. To induce senescence, cells were passaged up to 25 times with a CD144 enrichment step performed after every 4^th^ passage using anti-CD144 magnetic coated beads. To delete CSE *in vitro*, cells (passage 3) were treated with 4-OH-tamoxifen (10 μmol/L, Sigma, Darmstadt, Germany), after 5 days the 4-OH-tamoxifen was removed and the cells were further passaged for 2-3 times before experiments were performed. Cells isolated from wild-type littermate mice were treated identically.

For studies involving supplementation of the ATP, cells were treated with ATP-polyamine-biotin (20 mmol/L, # HY-D0183-1mg, CliniSciences, Nanteree, France), for 24 hours prior to harvesting.

For studies involving treatment with doxorubicin, cells were treated with doxorubicin (1 μmol/L, Sigma) for 24 hours prior to harvesting.

**Replicative senescence (multiduplication-induced senescence).**

Cells were retrieved from cryopreservation and 4000 cells per condition were seeded in 48 well plates, every 48 hours cells were disassociated (accutase), counted using a cell counter (CASY, cell counter, OMNI Life Science GmbH, Bremen, Germany)and were plated in coated (as described above) 48 well plates in a density of 4000 cells per well. Every cycle of disassociation is referred to as a passaging cycle in the figure legends and the text.

**Endothelial cell proliferation *in vitro***

Endothelial cells were seeded onto fibronectin-coated 24 well plates at a density of 5000 cells per well. Cells were allowed to adhere for 12 hours in the absence of serum in ECGM-2 medium (Lonza, Cologne, Germany) to achieve synchronization. Thereafter, cells were maintained for up to 120 hours before being disassociated from the culture plates and counted using a cell counter (CASY, cell counter, OMNI Life Science GmbH, Bremen, Germany).

**Cell cycle analyses**

Endothelial cells were seeded onto fibronectin-coated 6 well plates at a density of 150000 cells per well and allowed to adhere for 12 hours in the absence of serum in ECGM-2 medium (Lonza, Cologne, Germany) to achieve synchronization. Thereafter, cells were maintained for an additional 24 hours in serum and growth factor enriched culture media. Experiments were stopped by the addition of ice-cold PBS and single-cell suspensions were generated using accutase solution (Sigma, Darmstadt, Germany), and cells were recovered by centrifugation (300 g, 4 minutes). Cell pellets were then gently resuspended in 0.3 mL PBS containing serum (50%), before being fixed by the dropwise addition of 0.9 mL of ice-cold ethanol (70%). Samples were then incubated for 2 hours at 4 °C before being washed twice with cold PBS/azide to remove ethanol and precipitated protein. Propidium iodide (50 μg/mL in PBS containing 100 U/ml RNAse) was added and samples were incubated for at least 30 minutes at room temperature and propidium iodide incorporation was assessed using a FACSVerse instrument and analyzed with the FACSflow software (BD, Heidelberg, Germany).

**Senescence associated β galactosidase (SAβG) activity**

Two methods were used to evaluate SAβG activity. The Beta-Glo assay system (# E4720, Promega GmbH, Waldorff, Germany) was used to assess SAβG in native human endothelial cells immediately after FACs sorting, according to the manufacturer’s instructions. In brief, 100.000 endothelial cells were incubated in 100 µL EGM2 media (Lonza, Cologne, Germany) supplemented with 5% orthologous human serum and 100 µL Beta Glo reagent (1:1 ratio of β-galactosidase substrate and buffer) and mixed gently at room temperature for 30 seconds. After 30 minutes at room temperature the signal was assessed using a luminometer (Spark, Tecan Group Ltd, Männedorf, Switzerland) with an integration time of 1 second.

For cultured murine endothelial cells SAβG was assessed using a Cellular Senescence assay kit (# KAA002, Merck, Darmstadt, Germany) according to the manufacturer’s instructions. In brief, cells were seeded onto fibronectin coated µ-Slide 8 Well slides (# 80826, ibidi GmbH, Gräfelfing, Germany) and cultured to confluence. Thereafter, cells were incubated with fixing solution (Part No. 2004755, 15 minutes, room temperature) and after washing 3 times with PBS, cells were incubated with 150 µL of X-gal solution (pH 6.0, buffered with 2N HCl) for 4 hours (37 °C). Where indicated in the text, additional antibody incubationwas performed on top of the SAβG signal. Cells were visualized in phase contrast mode using a Zeiss Axio Observer microscope (Zeiss, Oberkochen, Germany) and analyzed with the ZEN software (Zeiss, Oberkochen, Germany).

**Metabolomics**

For the evaluation of the metabolites involved in glycolysis, the pentose phosphate pathway and Krebs cycle, frozen cell samples were extracted with ice-cold methanol/water (85/15, v/v). After 10 minutes on ice with repeated vortexing, samples were centrifuged (11000 g, 4 °C, 10 minutes), and the supernatants were immediately processed for analysis. Isotope labeled internal standards were added and samples were evaporated in a vacuum concentrator (Eppendorf, Hamburg, Germany) at 30 °C. The samples were resolved in 50 μL water methanol/water (50/50, v/v) and subsequently transferred to the LC-MS/MS system. Liquid chromatography was performed on an Agilent 1290 Infinity pump system (Agilent, Waldbronn, Germany) with a Phenomenex Luna Amino-column (100 mm × 2.0 mm, 3 µm) with ammonium acetate (10 mmol/L, pH 9.0) as mobile phase A and 100 % acetonitrile as mobile phase B. Five µL per sample were injected. The column temperature was set at 30 °C. The gradient with a flow rate of 700 µL/ minute was as follows: 0-1 minutes, 5 % A; 1-3 minutes, 5-60% A; 3-15 minutes, 60-95 % A; 15-18 minutes, 95 % A; 18-18.1 minutes, 95-5 % A; 18.1-24.1 minutes, 5 % A. Mass spectrometry was performed using a QTrap 5500 mass spectrometer (Sciex, Darmstadt, Germany) with electrospray ionization in negative mode. ESI parameters were set to TEM 400 °C, IS -4500 V, CUR 25 psi, GS1 40 psi, and GS2 60 psi.

For the detection of free amino acids sample preparation was performed using the EZ:faast LC MS free amino acid analysis kit (Phenomenex, Aschaffenburg, Germany) according to the manufacturer’s instructions, with minor modifications. Internal standards were applied to all samples and to the standard curve. Sample pH was adjusted to between pH 1.5-6.0 with HCl. Metabolite analysis was performed by LC-MS/MS using the EZ:faast AAA-MS HPLC column (inner diameter 2 mm) on an Agilent 1290 Infinity LC system (Agilent, Waldbronn, Germany) coupled to a QTrap 5500 mass spectrometer (Sciex, Darmstadt, Germany). Electro spray ionization in positive mode was employed. Data acquisition and instrument control were managed through the software Analyst 1.6.2. Peak integration, data processing and quantification was performed using MultiQuant 3.0 (both Sciex, Darmstadt, Germany). Area under the peak was used for quantification of the metabolites and the specific MRM transitions were normalized to the appropriate isotope-labelled internal standards and to the protein content of the sample. Statistical analysis, pathway enrichment analysis and heat maps were generated with Metaboanalyst 5.0.^2^

**Animals**

Floxed CSE (CSE^fl/fl^) mice,^3^ were crossed with tamoxifen-inducible Cdh5-CreERT2 mice,^2^ to generate animals specifically lacking CSE in endothelial cells following treatment with tamoxifen as described.^4,5^ Mice were housed in conditions that conform to the Guide for the Care and Use of Laboratory Animals published by the U.S. National Institutes of Health (NIH publication no. 85-23). Animals received the usual laboratory diet and all studies were approved by the animal research ethic committee in Darmstadt (FU 1189 and FU1250). Littermates of both genders were used. To induce robust Cre activity, animals were treated with tamoxifen (75 mg/kg i.p., Sigma, Darmstadt, Germany) for 5 days. Knockdown of CSE was observed 7 days post-injection. For all animal studies, the allocation of animals to the different treatment groups was blinded to the investigator and was revealed only after the data had been analyzed. Animals were randomized using the block randomization method to ensure similar sample sizes per age and sex matched group. Experiments were performed in a double blinded manner and no animals were excluded from the analysis. In some experiments, adeno-associated viruses i.e., AAV9-CSE S377A and AAV9-CSE-S377D, were injected into the tail vein of 3 month old mice at a concentration of 10^12^ vg/mL (in 100 μL). The mice were then maintained under standard conditions for up to 30 months.

**Aortic regeneration model and endothelial proliferation *in vivo***

Abdominal aortic injury was performed as previously described.^6^ To visualize endothelial proliferation mice were injected 4 hours prior to sacrifice with 5-ethynyl-2 ́-deoxyuridine (Click-iT EdU; 50 mg/kg i.p.; Thermo Fisher Scientific, Dreieich, Germany). Aortae were prepared by removing the surrounding tissue and fixed with 4% paraformaldehyde for 2 hours at room temperature and washed with blocking buffer containing 3% BSA and 0.5% Triton X-100 in PBS. A Click-iT reaction cocktail containing Alexa Fluor 647 azide was prepared according to the manufacturer’s protocol. Some aortas were used for endothelial cell isolation and FACs sorting as described above.

**AAV9- SLRSPPS viral particle generation**

An adeno-associated (AAV)-based delivery system was used to overexpress Myc-tagged human CTH (CSE) and the indicated serine 377 to alanine (SA mutant) and serine 377 to aspartate (SD mutant) mutants in the murine endothelial cells *in vivo*. Myc-CSE or its variants were put under control of the human cytomegalovirus promoter (CMV) and linked to the fluorophore mCherry using a self-cleaving porcine teschovirus-1 2A (P2A) peptide by cloning the ORF into the AAV genome plasmid pSSV9-CMV-P2A-mCherry via AgeI and Bsu36I restriction sites. A CMV-mCherry construct in the same context served as control. As it has been shown to increase transduction efficiency in endothelial cells, an AAV9 capsid variant with the targeting peptide SLRSPPS was used for vector production.^7^ AAV vectors were generated by co-transfection of the respective pSSV9 genome plasmid, the helper plasmid p5E18VD2/9-SLRSPPS providing the AAV9-SLRSPPS capsid variant, and the adenoviral helper plasmid pDGΔVP into low passage HEK293T cells. The assembled recombinant AAV9 vectors were harvested from cell culture supernatant and cell lysates, purified by discontinuous iodixanol gradient ultracentrifugation, and titrated via quantitative real-time PCR.^8^ AAV vectors were intravenously injected via the tail vein of 3 month old mice at a concentration of 10^12^ vg/mL (in 100 μL).

**Telomere length**

Telomere length was determined using a monochrome multiplex (MM) qPCR method as described.^9^ In brief, genomic DNA was isolated from endothelial cells with the DNeasy Blood and Tissue Kit (# 69504, Qiagen, Hilden, Germany) and preserved in Tris-HCl (10 mmol/L, pH 7.5) containing EDTA (0.1 mmol/L), at a concentration of 100 ng/µL. For MM-qPCR, 5 ng genomic DNA was used per reaction. Primer pairs used for telomere amplification were Telg: 5′-ACACTAAGGTTTGGGTTTGGGTTTGGGTTTGGGTTAGTGT-3′ and Telc: 5′–TGTTAGGTATCCCTATCCCTATCCCTATCCCTATCCCTAACA–3′. For reference, a signal for human β-globin was acquired at 88 °C using the primers hbgu 5′–CGGCGGCGGGCGGCCGGGGCTGGGCGGCTTCATCCACGTTCACCTTG–3′ and hbgd 5′–GCCCGGCCCGCCGCGCCCGTCCCGCCGGAGGAGAAGTCTGCCGTT–3′. All measurements were carried out in triplicate in a single-blinded manner. Telomere length was calculated as the ratio of the number of copies of the telomere template (T) versus the β-globin template (S) and normalized to the values obtained for young or wild-type mice.

**Droplet digital telomeric repeat amplification protocol (ddTRAP)**

To measure telomerase activity, the QX200 Droplet Digital PCR system (Bio-Rad Laboratories, Hercules, CA, USA) system was used according to the manufacturer’s instructions and as previously described.^10^ The extension reaction was performed with a TS telomerase extension substrate (HPLC purified, 5´‐AAT CCG TCG AGC AGA GTT‐3’) for 30 minutes at 37 °C in TRAP buffer with 25,000-50,000 endothelial cells. Afterwards, 200 cell equivalents of the extension reaction were used for the ddPCR reaction. QX200 ddPCR EvaGreen supermix (Bio‐Rad, 186‐4034) was used with 250 nmol/L of TS and 250 nmol/L of ACX primers (TS: 5´-AAT CCG TCG AGC AGA GTT-3´, ACX: 5´-GCG CGG CTT ACC CTT ACC CTT ACC CTA ACC-3´). After droplet generation, the PCR reaction was carried out in a C1000 Touch Thermal Cycler (Bio-Rad Laboratories, Hercules, CA, USA) using the following conditions: 95 °C for 5 minutes, 45 cycles of 95 °C at 30 seconds), 53 °C for 20 seconds and 72 °C for 30 seconds. Signals were measured with a QX200 Droplet Reader (Bio-Rad Laboratories, Hercules, CA, USA) using the QuantaSoft Reading software in the absolute quantification mode (Bio‐Rad, Version 1.74). Analysis was performed with QuantaSoft AnalysisPro software (Bio‐Rad) and all data shown were calculated above the threshold. All experiments included a no template control.

**Telomer Q-fluorescent in situ hybridization (FISH)**

Telomere Q-FISH was performed as previously described with modifications.^11^ Endothelial cells were cultured until confluency in fibronectin coated dishes and fixed with methanol/acetic acid. Cells were incubated with PBS for 15 minutes prior to fixation in 4% formaldehyde in PBS for 4 minutes at 37 ^°^C. Murine aortas were fixed immediately after dissection in 4% formaldehyde in PBS for 15 minutes at 37 °C. Samples were washed with PBS (3 x 5 minutes, 37 °C) and treated with RNase A solution (100 μg/mL in 2 x saline-sodium citrate [SSC] buffer containing 0.3 mol/L NaCl and 30 mmol/L sodium citrate, pH 7.0) for 1 hour at 37 ^°^C. After washing in 2x SSC buffer (3 x 5 minutes, 37 °C) samples were incubated with pepsin (0.005% in 0.01 mol/L HCl, pH 2.0) for 4 minutes at 37°C. Thereafter, cells were washed with PBS (3 x 5 minutes, 21 °C) and dehydrated in cold ethanol series (1 minute in 70%, 85%, 100%). Samples were heated to 80 °C for 5 minutes and incubated with 100 µL PNA probes (200 nmol/L), at 85 °C for 10 minutes to achieve denaturation. The PNA Fish probes used were Cy3 labeled, C-rich telomere probe (# F1002, PNA Bio, CA, USA) and Cy5 labeled centromere probe (# F3005, PNA Bio, CA, USA). Samples were kept in the dark at room temperature for additional 1 hour before being washed with 2 x SSC/0.1% Tween-20 (3 x 10 minutes, 55-60 °C). DAPI staining was performed for 15 minutes at room temperature with 2 x SSC buffer containing DAPI (20 ng/mL). Finally, after washing with 2 x SSC, SSC and distilled water, slides were mounted with a DAKO fluorescence mounting media and visualized using a Zeiss LSM780 microscope (Zeiss, Oberkochen, Germany).

**RNA sequencing, ATAC sequencing and analysis**

To extract total RNA from endothelial cells, samples were processed with the RNeasy Mini Kit (Qiagen, Hilden, Germany) and subjected to on-column DNase digestion according to the manufacturer’s protocol. RNA and library preparation integrity were verified with LabChip Gx Touch 24 (Perkin Elmer). 4 µg of total RNA was used as input for VAHTS Stranded mRNA-seq Library preparation following manufacture’s protocol (Vazyme). Sequencing was performed on NextSeq2000 instrument (Illumina) using P3 flowcell with 1x72bp single end setup. The resulting raw reads were assessed for quality, adapter content and duplication rates with FastQC (http://www.bioinformatics.babraham.ac.uk/projects/fastqc). Trimmomatic version 0.39 was employed to trim reads after a quality drop below a mean of Q20 in a window of 5 nucleotides.^12^  Only reads between 30 and 150 nucleotides were cleared for further analyses. Trimmed and filtered reads were aligned versus the Ensembl human genome version hg38 (Ensembl release 104) using STAR 2.7.9a with the parameter “--outFilterMismatchNoverLmax 0.1” to increase the maximum ratio of mismatches to mapped length to 10%.^13^ The number of reads aligning to genes was counted with featureCounts 2.0.2 tool from the Subread package.^14^ Only reads mapping at least partially inside exons were admitted and aggregated per gene. Reads overlapping multiple genes or aligning to multiple regions were excluded. Differentially expressed genes were identified using DESeq2 version 1.30.0.^15^ Only genes with a minimum fold change of ± 2 (log2 = ±1), a maximum Benjamini-Hochberg corrected *p*-value of 0.05, and a minimum combined mean of 5 reads were deemed to be significantly differentially expressed. The Ensemble annotation was enriched with UniProt data (release 06.06.2014) based on Ensembl gene identifiers (Activities at the Universal Protein Resource (UniProt)).

For ATAC sequencing cryopreserved human native cells were immediately processed. In brief, 50,000 cells were centrifuged at 500 g for 5 min at 4 °C and washed with PBS. The Cell pellet was resuspended in 50 µl lysis/transposition reaction (12.5 µl THS-TD-buffer, 2.5 µl Tn5, 5 µl 0.1% digitonin and 30 µl water) and incubated at 37 °C for 30 min with occasional snap mixing. Following purification of the DNA fragments was done by MinElute PCR Purification kit (Qiagen). Amplification of Library together with Indexing Primers was performed as described elsewhere (transposition of native chromatin for fast and sensitive epigenomic profiling of open chromatin, DNA-binding proteins and nucleosome position.^16^ Libraries were mixed in equimolar ratios and sequenced on NextSeq2000 platform using 2x36bp paired-end setup.

Raw reads were trimmed according to RNA-seq reads and aligned versus the human genome version hg38 (Ensembl release 104) using STAR 2.7.10a with the parameters “--outFilterMismatchNoverLmax 0.1  --outFilterMatchNmin 20 --alignIntronMax 1 --alignSJDBoverhangMin 999 --outFilterMultimapNmax 1 --alignEndsProtrude 10 ConcordantPair” ^13^ and retaining only unique alignments to exclude reads of uncertain origin. Reads were further deduplicated using Picard 2.25.5 to mitigate PCR artefacts leading to multiple copies of the same original fragment. Reads aligning to the mitochondrial chromosome were removed. The Macs2 peak caller version 2.1.1 was employed to accommodate for the range of peak widths typically expected for ATAC-Seq.^17^ Minimum qvalue was set to -4 and FDR was changed to 0.0001. Peaks overlapping ENCODE blacklisted regions (known misassemblies, satellite repeats) were excluded.

In order to be able to compare peaks in different samples to assess reproducibility, the resulting lists of significant peaks were overlapped and unified to represent identical regions. Sample counts for union peaks were produced using bigWigAverageOverBed (UCSC Toolkit) and normalized with DESeq2 1.30.0 to compensate for differences in sequencing depth, library composition, and ATAC-Seq efficiency.^18^ Peaks were annotated with the promoter of the nearest gene in range (TSS ± 5000 nt) based on reference data of GENCODE vM15 and the UROPA tool.^19^

TF binding analysis and TFBS prediction for TP53 was performed by a digital genomics footprinting approach and the TOBIAS tool.^20^ Thereby, footprints were detected and scored (scoreBigWig module) after Tn5 signal correction (ATAC_correct module), and differentially bound sites were predicted and reported (Bindetect module). TP53 targets were extracted from all potential TFBS binding sites according to the TOBIAS scoring model.

**H_2_S_n_ measurements**

Snap frozen endothelial cells were used and the specific products of the reaction of H_2_S with sulfane sulfur probe 4 (SSP4; Dojindo, GERBU Biotechnik GmbH, Heidelberg, Germany) were quantified by LC-MS/MS as described.^21^

**Real time-quantitative PCR (RT-qPCR)**

Total RNA was extracted using TriReagent (Merck) according to the manufacturer’s protocol. For the generation of cDNA, total RNA was reverse transcribed using the SuperScriptIII (Life Technologies GmbH, Darmstadt, Germany) and random hexamer primers (Promega, Madison, USA). Messenger RNA levels were quantified using the cycle threshold (C_T_) values determined by SYBR green qPCR master mix (SensiFAST SYBR Lo-ROX Bioline, London, UK) in a MIC qPCR cycler (BMS, Upper Coomera, Australia). Messenger RNA levels were normalized to 18S RNA. The following primers were used for the qPCR:

| Species | Gene | Sequence |
| --- | --- | --- |
| Mouse | TERT Fw | 5’-CGTTCCTGTTCTGGCTGATG-3’ |
| Mouse | TERT Rev | 5’-TGATGCCTGACCTCCTCTTG-3’ |
| Mouse | TERC Fw | 5’-AACAAACGTCAGCGCAGGAG-3’ |
| Mouse | TERC Rev | 5’-TCAGGTAAGACACCGAACAC-3’ |
| Human | P21 Fw | 5’-TGGAGACTCTCAGGGTCGAAA-3’ |
| Human | P21 Rev | 5’-GGCGTTTGGAGTGGTAGAAATC-3’ |
| Human | P53 Fw | 5’-GAGGTTGGCTCTGACTGTACC-3’ |
| Human | P53 Rev | 5’-TCCGTCCCAGTAGATTACCAC-3’ |
| Human | P16 Fw | 5’-ACCAGAGGCAGTAACCATGC-3’ |
| Human | P16 Rev | 5’-TGTCGTTCGCGGGCGCAACTG-3’ |
| Mouse | p21 Fw | 5’-CGAGAACGGTGGAACTTTGAC-3’ |
| Mouse | p21 Rev | 5’-CAGGGCTCAGGTAGACCTTG-3’ |
| Mouse | p53 Fw | 5’-GTCACAGCACATGACGGAGG-3’ |
| Mouse | p53 Rev | 5’-TCTTCCAGATGCTCGGGATAC-3’ |
| Mouse | p21 Fw | 5’-CGAGAACGGTGGAACTTTGAC-3’ |
| Mouse | p21 Rev | 5’-CAGGGCTCAGGTAGACCTTG-3’ |
| Mouse | p16 Fw | 5’-GAACTCTTTCGGTCGTACCC-3’ |
| Mouse | p16 Rev | 5’-TGGGCGTGCTTGAGCTGA-3’ |
| Human/Mouse | 18S Fw | 5’-CTTTGGTCGCTCGCTCCTC-3’ |
| Human/Mouse | 18S Rev | 5’-CTGACCGGGTTGGTTTTGAT’-3’ |

**Adenoviral transduction**

GFP and CSE adenoviruses were generated as described.^22,23^ Adenoviruses (10 MOI) were incubated with AdenoBoost (Sirion Biotech, Martinsried, Germany) for 30 minutes before being added to endothelial cells cultured in DMEM/F12 containing 0.1% bovine serum albumin (BSA) for 4 hours at 37 °C. The cells were then washed, the medium were replaced and cells were cultured for the times indicated in the results section i.e. for human native endothelial cells up to 6 days, for murine CSE^iΔEC^ endothelial cells from passage 4 up to passage 7 and for wild-type cells from passage 4 up to passage 25. The latter cells went through 3 rounds of transduction i.e. every 6 passages, before use. In some experiments, GFP+ cells were isolated (FACS sorting) for further evaluation of telomerase activity and protein expression.

**Lentiviral generation and transduction**

Guide RNAs against human TP53 (antisense: AAACTCGCTATCTGAGCAGCGCTCC and sense: CACCGGAGCGCTGCTCAGATAGCGA) and TERT (antisense: AAACCTACGGGGTGCTCCTCAAGAC and sense: CACCGTCTTGAGGAGCACCCCGTAG) were designed with an online tool (<http://chopchop.cbu.uib.no/>). A non-targeted control was also used (antisense: AAACAGGACTTGTTAGCCCGGAAC and sense: CACCGTTCCGGGCTAACAAGTCCT). For overexpression plasmids p53WT and p53K120R mutant, where cloned into a TK-PCDH-copGFP-T2A-puro vector (Genomeditech, Shanghai, China). For CRIPSR-Cas9 genome editing, annealed gene specific gRNAs were cloned into a plentiCRISPRv2 plasmid co-expressing a Cas9 nuclease and a puromomycin-selection marker (Addgene, 98290). Lentivirus production was performed by co-transfection of 80% confluent HEK293FT cells (cultured in DMEM/F-12, Gibco supplemented with 10% FCS and 1% pen-strep) with pMD2.G (Addgene, 12259), psPAX2 (Addgene, 12260) and the target plasmids in OptiMEM in the presence of polyethylamine (1 µg/µl, Sigma). Supernatant containing viral particles were collected 72 and 96 hours after transfection, filtered through a 0.45μm filter and stored in -80 ^o^C until use.

Human native endothelial cells were transduced for 24 hours with 1:1 ECGM-2: Viral volumes in the presence of 8 µg/ml polybrene (Santa Cruz). After transduction, cells were cultured for 24 hours with ECGM-2 and selected with ECGM-2 media containing 1 µg/ml puromycin (Invivogen). Selected cells were allowed to recover for additional 48 hours in ECGM-2 prior to other experiments.

**Small interfering RNA**

Murine lung endothelial cells were treated with control siRNA (25 pmol) or siRNA directed against Glut1 (25 pmol, # NM_011400, Merck, Darmstadt, Germany) or p53 (25 pmol, # Cell Signalling Technologies, Frankfurt, Germany) and human native endothelial cells were treated with control siRNA (25 pmol) or siRNA directed against Glut1 (25 pmol, # 117394, Thermo Fischer Scientific, Dreieich, Germany) for 72 hours prior to experimentation. Transfection was performed in OptiMEM media (# 11058021, Thermo Fischer Scientific, Dreieich, Germany) using Lipofectamine RNAi Max (# 13778030, Invitrogen, Darmstadt, Germany) according to the manufacturer’s instructions.

**Nuclear and cytosolic extraction**

Endothelial cells (~500 000) were washed with PBS and immediately collected in 1.5 mL tubes and recovered by centrifugation (200 g, 21 °C, 5 minutes). The supernatant was removed and the pellet was resuspended in 150 μL of cytoplasmic extract (CE) buffer (0.075% v/v NP-40 in 10 mmol/L HEPES, 60 mmol/L KCl, 1 mmol/L EDTA, 1 mmol/L DTT and 1 mmol/L PMSF, adjusted to pH 7.6). Samples were incubated on ice for 5 minutes and recovered by centrifugation (1000 g, 10 minutes). The supernatant containing the cytoplasmic extract was transferred in a new tube and centrifuged again (16.000 g. 4 °C, 10 minutes) and the resulting supernatant was stored until further analyzed. The pellet containing nuclei was washed 3 times with 1 mL of CE buffer without NP-40. Following the last washing step pelleted nuclei were resuspended in 50 μL of nuclear extraction buffer (20 mmol/L Tris HCl, 420 mmol/L NaCl, 1.5 mmol/L MgCl_2_, 0.2 mmol/L EDTA, 1 mmol/L PMSF and 25% (v/v) glycerol, adjusted to pH 8.0) and mixed by vortexing. After incubation on ice for 10 minutes and periodical vortexing, samples were centrifuged (16.000 g, 4 °C, 10 minutes) and the supernatant was stored for further analysis.

**Immunoprecipitation**

Immunoprecipitations were performed using antibodies directed against p53, GST-CSE and Tert. In the case of *in vitro* pull down studies, the interaction of GST-CSE (5 µg, purified as described^24^) and recombinant p53 protein (5 µg, # ab82201, Abcam, Berlin, Germany) was assessed using glutathione magnetic agarose beads (# 78602, Thermo Fischer Scientific, Dreieich, Germany). For immunoprecipitation of endogenous proteins, total protein (500 µg) was incubated with anti-p53 or anti-TERT antibodies (1µg/sample) pre-coated on sepharose G beads. Following overnight incubation, proteins were recovered and eluted in a buffer containing 3% SDS, 1% β-mercaptoethanol and 0.005% bromophenol blue in PBS (15 minutes, 95 ^o^C) and detected with SDS-PAGE followed by immunoblotting.

To compare the interaction of p53 with the wild type and mutated CSE, Myc tagged constructs with a serine to alanine (SA) or serine to aspartate (SD) mutation at position 377 were used as previously described ^4^. HEK239 cells were transfected with Lipofectamine 2000 transfection reagent (# 11668030, Thermo Fisher Scientific, Dreieich, Germany) according to the manufacturer’s instructions. Thirty six hours later cells were lysed with ice-cold buffer (50 mmol/L Tris HCl-pH 7.5, 150 mmol/L NaCl, 25 mmol/L NaF,10 mmol/L Na_4_P_2_O_7_,1% Triton X-100 and 0.5% sodium deoxycholate) and 500 µg of total protein was used for Myc immunoprecipitation with the use of the agarose pre-coated beads (# sc-40 AC, Santa Cruz Biotechnology (Heidelberg, Germany), and detected with SDS-PAGE followed by immunoblotting of p53.

**Immunoblotting**

Samples were lysed in ice-cold RIPA buffer (50 mmol/L Tris HCl-pH 7.5, 150 mmol/L NaCl, 25 mmol/L NaF,10 mmol/L Na_4_P_2_O_7_,1% Triton X-100 and 0.5% sodium deoxycholate) supplemented with 0.1% SDS and protease and phosphatase inhibitors. Protein concentrations were determined using the Bradford assay, and detergent-soluble proteins were solubilized in SDS sample buffer, separated by SDS-PAGE and subjected to Western blotting as described ^1^. Proteins were visualized by enhanced chemiluminescence using a commercially available kit (Amersham, Freiburg, Germany).

**Proximity ligation assay**

Proximity ligation was performed using the Duolink assay according to manufacturer’s instructions (MilliporeSigma, Darmstadt, Germany), using antibodies against p53 (mouse), CSE (rabbit), MOZ (rabbit), TERT (rabbit) or Hsp9a (mouse). Images were taken using a confocal microscope (LSM-780; Zeiss, Oberkochen, Germany) and ZEN software (Zeiss, Oberkochen, Germany).

**Immunochistochemistry**

Endothelial cells and murine aortae were fixed with 4% paraformaldehyde for 15 minutes at room temperature prior to staining. Human arteries were fixed with 4% paraformaldehyde for 1 hour at room temperature, embedded in OCT Tissue TeK (Sakura, Staufen, Germany) and frozen on dry ice. Thereafter, sections (6 μm) were prepared on a microtome (Microm HM 650, Thermo Scientific, Darmstadt, Germany) and placed on positively charged glass slides. The slides were washed with PBS for 5 minutes and incubated 2 hours at room temperature (RT) in a blocking buffer consisting of with Triton X-100 (0.3%), donkey serum (5%) and BSA (0.5%) in PBS. Samples were washed with PBS and incubated overnight (4°C) with primary antibodies in PBS containing Triton X-100 (0.2%). Thereafter, secondary antibodies against rabbit and rat and mouse were diluted 1:200 in PBS supplemented with DAPI (200 ng/mL). After washing, sections were mounted with Dako fluorescent mounting medium (Dako, Glostrup, Denmark). Images were taken using a confocal microscope (LSM-780; Zeiss, Oberkochen, Germany) and ZEN software (Zeiss, Oberkochen, Germany). Co-localization signals were exported from the ZEN software.

**Chromatin immunoprecipitation**

Cell extracts, and the crosslinking and isolation of nuclei was performed with the truCHIP™ Chromatin Shearing Kit (Covaris, USA) according to the manufacturer’s instructions. After sonification of the lysates (4 °C) for 10 cycles of 30 seconds on and 90 seconds off (Bioruptur Plus, Diagenode, Seraing, Belgium), cell debris was removed by centrifugation (10 000 g for 5 minutes at 4 °C) and the lysates were diluted 1:3 in dilution buffer (20 mmol/L Tris/HCl pH 7.4, 100 mmol/L NaCl, 2 mmol/L EDTA, 0.5% Triton X-100 and protease inhibitors). Pre-clearing was performed with 20 µL DiaMag protein A and protein G coated magnetic beads slurry (Diagenode, Seraing, Belgium) for 45 minutes at 4 °C. Thereafter, 5% of each sample was put aside and served to determine the input signal and the remainder was incubated as overnight (4 °C) with p53 antibody. p53 was then recovered following incubation with 50 µL DiaMag protein A and protein G coated magnetic beads (Diagenode, Seraing, Belgium) for 3 hours at 4 °C, and washed twice for 5 minutes with wash buffers 1-3 (wash buffer 1: 20 mmol/L Tris/HCl pH 7.4, 150 mmol/L NaCl, 0.1% SDS, 2 mmol/L EDTA, 1% Triton X-100; wash buffer 2: 20 mmol/L Tris/HCl pH 7.4, 500 mmol/L NaCl, 2 mmol/L EDTA, 1% Triton X-100; wash buffer 3: 10 mmol/L Tris/HCl pH 7.4, 250 mmol/L lithium chloride, 1% Nonidet p-40, 1% sodium deoxycholate, 1 mmol/L EDTA) and finally washed with TE-buffer pH 8.0. Elution of the beads was achieved using elution buffer (0.1 M NaHCO3, 1% SDS) containing 1x Proteinase K (Diagenode, Seraing, Belgium) and shaking at 600 rpm for 1 hour at 55 °C, followed by 1 hour at 62 °C and 10 minutes at 95 °C. After removal of the beads, the eluate was purified with the QiaQuick PCR purification kit (Qiagen, Hilden, Germany) and subjected to qPCR analysis. The following primers for the Glut1 and p21 promoters were used: Glut1, FW 5′-GGTTCTTTCTTCCACCGCGT-3′ and REV 5′-AGCAAGAATCCCAACCCCG-3′, and p21: FW 5′-GTGGCTCTGATTGGCTTTCTG-3′ and REV 5′-CTGAAAACAGGCAGCCCAAG-3′). Primers targeting the GAPDH promoter (FW: 5′-TGG TGT CAG GTT ATG CTG GGC CAG-3′ and REV 5′-GTG GGA TGG GAG GGT GCT GAA CAC-3′) were included as a negative control.

**In vitro assay of CSE activity**

The GST-CSE plasmid was generated and purified as described previously.^24^ In brief, 5 μg GST-CSE was allowed to react for 15 minutes at 37 ^o^C in the presence of PLP (10 nmol/L) and recombinant p53 (5 µg) before GST was immunopercipitated with glutathione beads. As a negative control GST-CSE pull down in the absence of p53 was performed. The activity of purified CSE (5 μg) in the absence and presence of p53, was assessed as described,^25^ in the absence and presence of L-cysteine, L-cystathionine (both 100 μmol/L), and PLP (10 nmol/L). As a negative control 5 µg of GST-CSE was denatured by heating to 100 °C for 10 minutes.

**p53 *S*-sulfhydration by antibody array**

An antibody directed against p53 (# 90001-MM06, SinoBiological, Eschborn, Germany; 10 ng/reaction) was added to a 96 well plate (3D-NHS Surface, PolyAn, Berlin) at a final volume of 50 μL in PBS buffer (150 mmol/L Na_2_HPO_4_ / NaHPO_4_ and 50 mmol/L NaCl, pH 8.5). BSA (50 µL, 5%) in TBST (137 mmol/L NaCl and 20 mmol/L Tris base, pH 7.4, supplemented with 0.1% Tween-20) and containing 0.002% NaN_3_ was used as a negative control. The plate was covered and incubated at overnight with agitation (4 °C). The solutions were discarded, and the wells were washed 5 times with 1.5 x PBS buffer supplemented with 0.01% Tween-20. All wells were then blocked with ethanolamine (50 mmol/L) in 100 mmol/L Tris at pH 9 for 2 hours at room temperature with agitation. The blocking solution was discarded, and wells were re-blocked with 5% BSA in TBS with 0.01% Tween-20 for a further 60 minutes. After washing, samples (0.5 µg protein/mL in a total volume of 100 µL) were added and incubated at 4 ^o^C overnight. After washing, the plate was recorded at 488 nm and 633 nm (Perkin Elmer, Ueberlingen, Germany). RIPA lysates of endothelial cells were labelled as described above.

**Glucose uptake**

Endothelial cells were seeded in in fibronectin coated µ-Slide 8 Well slides (#80826, ibidi GmbH, Gräfelfing, Germany) at 50% confluence and cultured for additional 24 hours. Thereafter, the culture medium was exchanged for medium lacking glucose for 1 hour prior to labelling. Murine aortas were collected in DMEM/F12 without glucose and allowed to stabilize in a humidified chamber for 2 hours prior to labelling. Samples were then supplemented with DMEM/F12 lacking glucose but supplemented with the fluorescent glucose analogue 2-(N-(7-nitrobenz-2-oxa-1,3-diazol-4-yl)amino)-2-deoxyglucose (2-NBDG, Merck, Darmstadt, Germany) at a final concentration of 100 µmol/L. After 30 minutes, cells/tissue were washed with PBS, fixed for 15 minutes at room temperature with 4% paraformaldehyde, co-stained with F-actin and visualized using a confocal microscope.

**^13^C-Glucose metabolic flux**

Endothelial cells were seeded on 10 cm culture dishes and allowed to grow to confluence before exchanging the culture medium for glucose-free medium supplemented with uniformly labelled ^13^C glucose (# 106032-62-6, Merck, Darmstadt, Germany) at a final concentration of 17 mmol/L. Thereafter, samples were collected and prepared as described.^26^ ^13^C-Labelled metabolites were detected by LC-MS/MS using an Agilent 1290 Infinity UPLC system (Agilent, Waldbronn, Germany) coupled to a QTrap5500 mass spectrometer (Sciex, Darmstadt, Germany), equipped with an ESI TurboIonSpray source. The flow rate was 0.22 mL/minute, autosampler temperature was set at 6 °C, and the column compartment was set to 35 °C. Separations were performed on an Asahipak NH2P-40 (250 × 2 mm, 4 μm particle size; Showa Denko, Munich, Germany). The mobile phase was composed of 20 mmol/L ammonium carbonate in H_2_O/5% acetonitrile (ACN), pH = 10 and ACN. After an initial 3.5 minute isocratic elution of 99.9% ACN, the percentage of ACN decreased to 85% at 3.6 minutes, to 75% at 8.1 minute, to 0% at 14 minutes, back to 99.9 % at 34 minutes. The composition was maintained at 99.9% ACN until 42 minutes. The QTrap5500-MS system was operated in triple quadrupole mode with positive/negative ion switching. An ion spray voltage of 5500/-4500 V was applied. Curtain gas was set to 30 psi, collision gas to medium, source temperature to 500 °C, ion source gas 1 to 35 psi, ion source gas 2 to 35 psi. Analyst 1.6.2 and MultiQuant 3.0 (both from Sciex, Darmstadt, Germany), were used for data acquisition and analysis.

**Glycolytic flux**

Endothelial cells (6 x 10^5^ cells per well) were seeded on a Seahorse XF96 culture plate pre-coated with fibronectin (Agilent, California, USA) six hours prior to measurements (XFe 96 extracellular flux analyzer, Agilent). Activators and inhibitors were purchased from Agilent and were used at the following concentrations: glucose (10 mmol/L) oligomycin (1.5 μmol/L), 2-deoxy-D-glucose (100 mmol/L), carbonyl cyanide 4-(trifluoromethoxy)phenylhydrazone (FCCP; 2.5 μmol/L), rotenone (0.5 μmol/L), antimycin A (0.5 μmol/L). An average of three technical replicates was measured for each cell batch.

**Telomerase interacting proteins**

Endothelial cells were seeded on 10 cm dishes precoated with fibronectin and grown to confluence. Subsequently, cells were harvested, and nuclear extracts were isolated as described above. Nuclear protein extract (100 µg) was used and 10 µL of sepharose fast flow protein A and G beads pre-coated with 1 µg anti-TERT antibody/ sample were added overnight (4 °C, with agitation). Following 3 washes with cytosolic extraction buffer without detergent, beads were resuspended in 50 µL 6 mol/L GdmCl, 50 mmol/L Tris/HCl (pH 8.5) and incubated at 95 °C for 5 minutes. Samples were diluted with 25 mmol/L Tris/HCl (pH 8.5) containing 10% acetonitrile to give a final GdmCl concentration of 0.6 mol/L. Proteins were digested with 1 µg trypsin (sequencing grade, Promega, Madison, USA) overnight (37 °C with gentle agitation). Digestion was stopped by adding trifluoroacetic acid to a final concentration of 0.5%. Peptides were loaded on multi-stop-and-go tips (StageTip) containing three strong cation exchange (SCX) disks and a stack of three C18-disks on top, and SCX fractionation by StageTips was performed as described.^27^ Three fractions of each sample were eluted in wells of microtiter plates and peptides were dried and resolved in 1% acetonitrile and 0.1 % formic acid. LC-MS/MS was performed using a Thermo Scientific Q Exactive Plus equipped with an ultra-high performance liquid chromatography unit (Thermo Scientific Dionex Ultimate 3000) and a Nanospray Flex Ion-Source (Thermo Scientific). Peptides were loaded on a C18 reversed-phase pre-column (Thermo Scientific) followed by separation on a 2.4 µm Reprosil C18 resin (Dr. Maisch GmbH) in-house packed picotip emitter tip (diameter 100 µm, 15 cm long from New Objectives, Littleton, USA) using a gradient from mobile phase A (4% acetonitrile, 0.1% formic acid) to a 50 % mobile phase B (99% acetonitrile, 0.1% formic acid) for 30 minutes. MS data were recorded by data dependent acquisition Top10 method for HCD fragmentation in positive mode. The full MS scan range was 300 to 2000 m/z with resolution of 70000, and an automatic gain control (AGC) value of 3 x E6 total ion counts with a maximal ion injection time of 160 ms. Only higher charged ions (2+) were selected for MS/MS scans with a resolution of 17500, an isolation window of 2 m/z and an automatic gain control value set to 10 x E5 ions with a maximal ion injection time of 150 ms. Fullscan data were acquired in profile and fragments in centroid mode. For data analysis MaxQuant 1.6.1. 0,^28^ Perseus 1.6.1.3 and Excel (Microsoft Office 2013) were used. For protein immunoprecipitation experiments the following settings were used: N-terminal acetylation (+42.01), oxidation of methionine (+15.99) and carbamidomethylation (+57.02) on cysteines were selected as modifications. The mouse reference proteome set (Uniprot, February 2018, 52538) and human reference proteome set (proteome identifier UP000005640) entries was used to identify peptides and proteins with a false discovery rate (FDR) less than 1%. Minimal ratio count for label-free quantification was 1. Reverse identifications and common contaminants were removed. Proteins were filtered to be at least identified 4 times in one experimental group. Missing values were replaced by random background values from normal distribution. Significant interacting proteins were determined by permutation-based FDR and student´s t-test.

**Statistics**

Data are expressed as mean ± SEM. Statistical evaluation was performed using Student's t test for unpaired data. One-way ANOVA (Bonferroni), two way ANOVA (followed by Tukey’s multiple comparisons test) were used where appropriate. Statistics for metabolomics were performed with Metaboanalyst 5.0. Statistical tests are described in the figure legend for each experiment. Values of *P*<0.05 were considered as statistically significant.

**Supplementary Tables**

**Supplementary Table 1.** **Ortho-PLSDA analysis of the most abundant metabolites in human native young and aged endothelial cells**

**Supplementary Table 2. Telomerase complexome identified in young and aged native human endothelial cells**

**Supplementary Table 3. Telomerase complexome identified in WT and CSE^iΔEC^ murine endothelial cells**

**Supplementary Table 4. Clinical data from the human subjects**

**References**

1.	Fleming I, Fisslthaler B, Dixit M, Busse R. Role of PECAM-1 in the shear-stress-induced activation of Akt and the endothelial nitric oxide synthase (eNOS) in endothelial cells. *J Cell Sci* 2005;**118:**4103–4111.

2.	Monvoisin A, Alva JA, Hofmann JJ, Zovein AC, Lane TF, Iruela-Arispe ML. VE-cadherin-CreERT2 transgenic mouse: a model for inducible recombination in the endothelium. *Dev Dyn* 2006;**235**:3413–3422.

3.	Syhr KMJ, Boosen M, Hohmann SW, Longen S, Köhler Y, Pfeilschifter J, Beck K-F, Geisslinger G, Schmidtko A, Kallenborn-Gerhardt W. The H_2_S-producing enzyme CSE is dispensable for the processing of inflammatory and neuropathic pain. *Brain Res* 2015;**1624**:380–389.

4.	Bibli S-I, Hu J, Sigala F, Wittig I, Heidler J, Zukunft S, Tsilimigras DI, Randriamboavonjy V, Wittig J, Kojonazarov B, Schürmann C, Siragusa M, Siuda D, Luck B, Abdel Malik R, Filis KA, Zografos G, Chen C, Wang DW, Pfeilschifter J, Brandes RP, Szabo C, Papapetropoulos A, Fleming I. Cystathionine γ lyase sulfhydrates the RNA binding protein human antigen R to preserve endothelial cell function and delay atherogenesis. *Circulation* 2019;**139**:101–114.

5.	Bibli S-I, Hu J, Looso M, Weigert A, Ratiu C, Wittig J, Drekolia MK, Tombor L, Randriamboavonjy V, Leisegang MS, Goymann P, Delgado Lagos F, Fisslthaler B, Zukunft S, Kyselova A, Justo AFO, Heidler J, Tsilimigras D, Brandes RP, Dimmeler S, Papapetropoulos A, Knapp S, Offermanns S, Wittig I, Nishimura SL, Sigala F, Fleming I. Mapping the endothelial cell S-sulfhydrome highlights the crucial role of integrin sulfhydration in vascular function. *Circulation* 2021;**143**:935–948.

6.	McDonald AI, Shirali AS, Aragón R, Ma F, Hernandez G, Vaughn DA, Mack JJ, Lim TY, Sunshine H, Zhao P, Kalinichenko V, Hai T, Pelegrini M, Ardehali R, Iruela-Arispe ML. Endothelial regeneration of large vessels is a biphasic process driven by local cells with distinct proliferative capacities. *Cell Stem Cell* 2018;**23**:210-225.e6.

7.	Varadi K, Michelfelder S, Korff T, Hecker M, Trepel M, Katus HA, Kleinschmidt JA, Müller OJ. Novel random peptide libraries displayed on AAV serotype 9 for selection of endothelial cell-directed gene transfer vectors. *Gene Ther* 2012;**19**:800–809.

8.	Jungmann A, Leuchs B, Rommelaere J, Katus HA, Müller OJ. Protocol for efficient generation and characterization of adeno-associated viral vectors. *Hum Gene Ther Methods* 2017;**28**:235–246.

9.	Cawthon RM. Telomere length measurement by a novel monochrome multiplex quantitative PCR method. *Nucleic Acids Res* 2009;**37**:e21.

10.	Leisegang MS, Bibli S-I, Günther S, Pflüger-Müller B, Oo JA, Höper C, Seredinski S, Yekelchyk M, Schmitz-Rixen T, Schürmann C, Hu J, Looso M, Sigala F, Boon RA, Fleming I, Brandes RP. Pleiotropic effects of laminar flow and statins depend on the Krüppel-like factor-induced lncRNA MANTIS. *Eur Heart J* 2019;**40**:2523–2533.

11.	Sacco A, Mourkioti F, Tran R, Choi J, Llewellyn M, Kraft P, Shkreli M, Delp S, Pomerantz JH, Artandi SE, Blau HM. Short telomeres and stem cell exhaustion model Duchenne muscular dystrophy in mdx/mTR mice. *Cell* 2010;**143**:1059–1071.

12.	Bolger AM, Lohse M, Usadel B. Trimmomatic: a flexible trimmer for Illumina sequence data. *Bioinformatics* 2014;**30**:2114–2120.

13.	Dobin A, Davis CA, Schlesinger F, Drenkow J, Zaleski C, Jha S, Batut P, Chaisson M, Gingeras TR. STAR: ultrafast universal RNA-seq aligner. *Bioinformatics* 2013;**29**:15–21.

14.	Liao Y, Smyth GK, Shi W. featureCounts: an efficient general purpose program for assigning sequence reads to genomic features. *Bioinformatics* 2014;**30**:923–930.

15.	Love MI, Huber W, Anders S. Moderated estimation of fold change and dispersion for RNA-seq data with DESeq2. *Genome Biol* 2014;**15**:550.

16.	Buenrostro JD, Giresi PG, Zaba LC, Chang HY, Greenleaf WJ. Transposition of native chromatin for fast and sensitive epigenomic profiling of open chromatin, DNA-binding proteins and nucleosome position. *Nat Meth* 2013;**10**:1213–1218.

17.	Zhang Y, Liu T, Meyer CA, Eeckhoute J, Johnson DS, Bernstein BE, Nusbaum C, Myers RM, Brown M, Li W, Liu XS. Model-based analysis of ChIP-Seq (MACS). *Genome Biol* 2008;**9**:R137.

18.	Anders S, Huber W. Differential expression analysis for sequence count data. *Genome Biol* 2010;**11**:R106.

19.	Kondili M, Fust A, Preussner J, Kuenne C, Braun T, Looso M. UROPA: a tool for Universal RObust Peak Annotation. *Sci Rep* 2017;**7**:2593.

20.	Bentsen M, Goymann P, Schultheis H, Klee K, Petrova A, Wiegandt R, Fust A, Preussner J, Kuenne C, Braun T, Kim J, Looso M. ATAC-seq footprinting unravels kinetics of transcription factor binding during zygotic genome activation. *Nat Communic* 2020;**11**:4267.

21.	Bibli S-I, Luck B, Zukunft S, Wittig J, Chen W, Xian M, Papapetropoulos A, Hu J, Fleming I. A selective and sensitive method for quantification of endogenous polysulfide production in biological samples. *Redox Biol* 2018;**18**:295–304.

22.	Bucci M, Papapetropoulos A, Vellecco V, Zhou Z, Pyriochou A, Roussos C, Roviezzo F, Brancaleone V, Cirino G. Hydrogen sulfide is an endogenous inhibitor of phosphodiesterase activity. *Arterioscler Thromb Vasc Biol* 2010;**30**:1998–2004.

23.	Coletta C, Papapetropoulos A, Erdelyi K, Olah G, Módis K, Panopoulos P, Asimakopoulou A, Gerö D, Sharina I, Martin E, Szabo C. Hydrogen sulfide and nitric oxide are mutually dependent in the regulation of angiogenesis and endothelium-dependent vasorelaxation. *Proc Natl Acad Sci USA* 2012;**109**:9161–9166.

24.	Asimakopoulou A, Panopoulos P, Chasapis CT, Coletta C, Zhou Z, Cirino G, Giannis A, Szabo C, Spyroulias GA, Papapetropoulos A. Selectivity of commonly used pharmacological inhibitors for cystathionine β synthase (CBS) and cystathionine γ lyase (CSE). *Br J Pharmacol* 2013;**169**:922–932.

25.	Stipanuk MH, Beck PW. Characterization of the enzymic capacity for cysteine desulphhydration in liver and kidney of the rat. *Biochem J* 1982;**206**:267–277.

26.	Grankvist N, Watrous JD, Lagerborg KA, Lyutvinskiy Y, Jain M, Nilsson R. Profiling the metabolism of human cells by deep 13C labeling. *Cell Chem Biol* 2018;**25**:1419-1427.e4.

27.	Rappsilber J, Mann M, Ishihama Y. Protocol for micro-purification, enrichment, pre-fractionation and storage of peptides for proteomics using StageTips. *Nat Protoc* 2007;**2**:1896–1906.

28.	Cox J, Mann M. MaxQuant enables high peptide identification rates, individualized p.p.b.-range mass accuracies and proteome-wide protein quantification. *Nat Biotechnol* 2008;**26**:1367–1372.

**Supplementary Table 4. Demographic data for the human subjects**

| **Characteristics** | **Young subjects** | **Aged subjects** |
| --- | --- | --- |
| No. | 50 | 50 |
| Mean age (range) | 20±3.4 | 80±2.3 |
| Male/Female | 36/14 | 39/11 |
| **Clinical data** |  |  |
| Hypertension | 0 | 5 |
| Diabetes | 0 | 0 |
| Hyperlipidemia | 0 | 5 |
| Coronary disease | 0 | 10 |
| Heart failure | 0 | 0 |
| Cancer | 12 | 16 |
