## Supplementary Figures for "A novel role for cystathionine γ lyase in the control of p53: impact on endothelial senescence and metabolic reprograming"

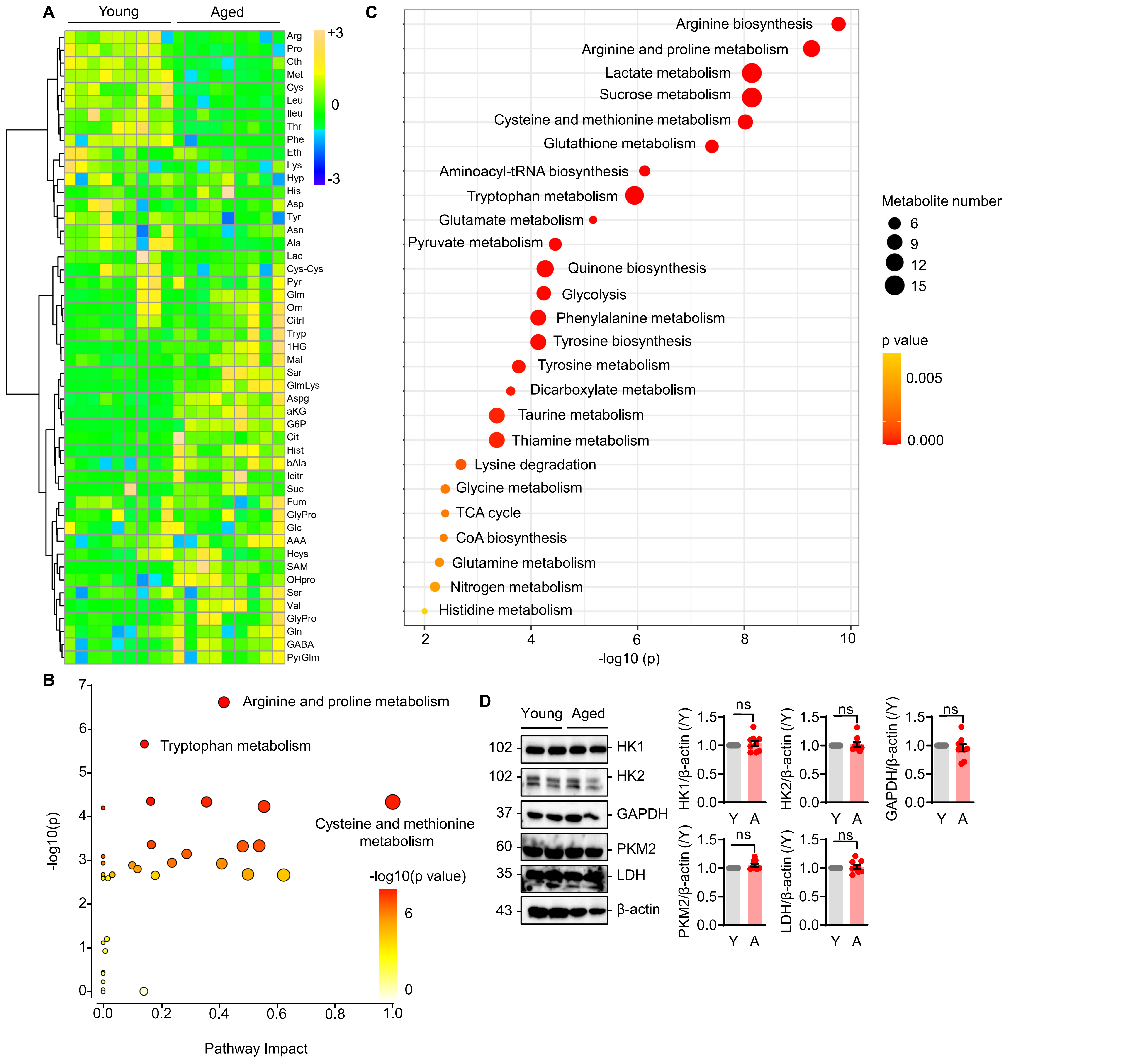


**Supplementary Fig. 1. Metabolomic profile of young and aged native human endothelial cells.** Targeted metabolomics analysis was performed using native mesenteric artery endothelial cells from young and aged individuals (each sample being a pool of cells from 5 different arteries); n=9/group. (**A**) Heatmap. (**B**) Metabolic pathway impact. (**C**) Enrichment ratio following pathway enrichment analysis. Statistical analyses performed with Metaboanalyst 5.0. (**D**) Representative immunoblotting and relative quantification of glycolytic enzymes in human endothelial cells from young and aged arteries; n=9/group, Student's t-test.

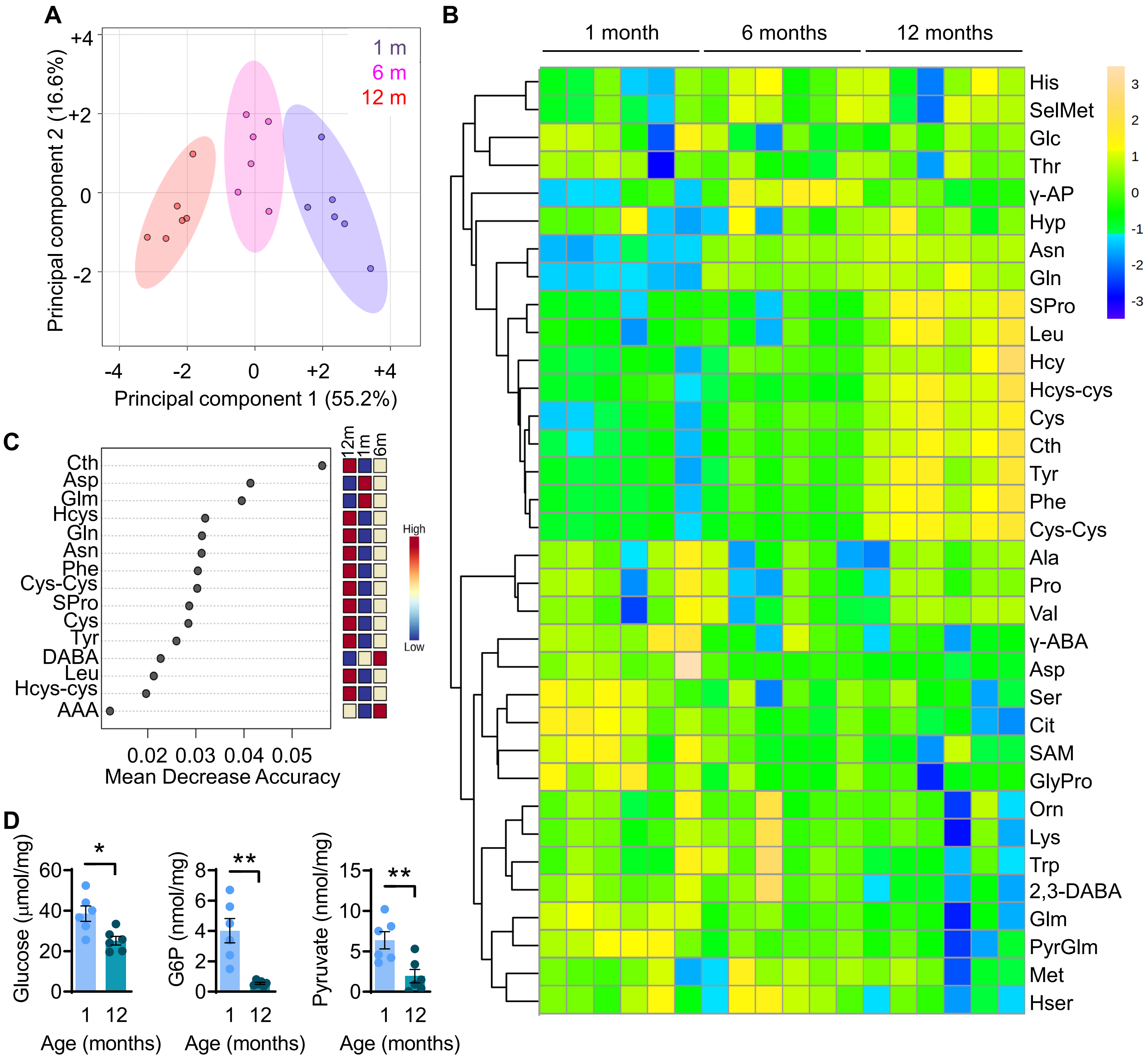


**Supplementary Fig. 2.** **Effects of aging on murine endothelial cell metabolism.** (**A**) Principal component analysis of the metabolic profile of aortic endothelial cells isolated from 1, 6 and 12 month old wild-type mice; n=6/group. (**B**) Heatmap of the most abundant metabolites from murine cells as in panel A; n=6/group. (**C**) Significant metabolites identified by Random Forest. The metabolites are ranked by the mean decrease in classification accuracy when they are permuted and show which ones primarily contribute to the metabolic difference among the tested groups. (**D**) Levels of glucose, glucose-6 phosphate (G6P) and pyruvate in aortic endothelial cells from 1 and 12 month old wild-type mice. *P<0.05, **P<0.01, Student's t-test.

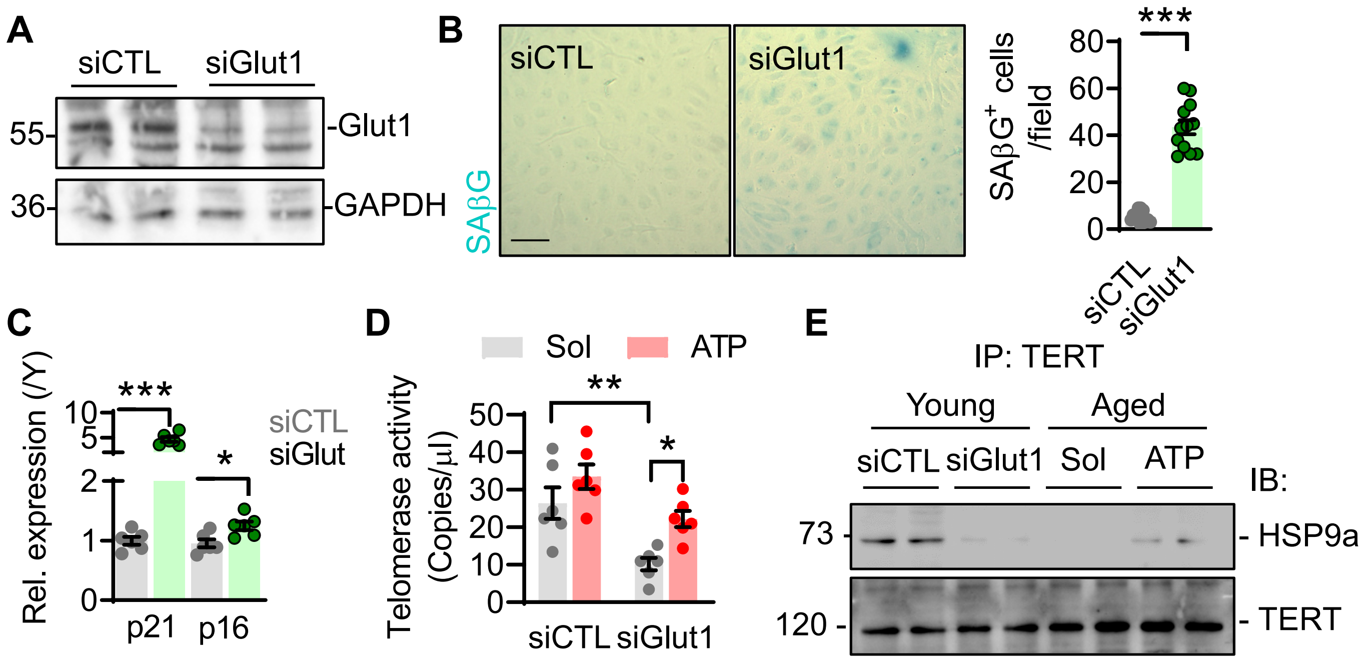


**Supplementary Fig. 3. Effects of Glut1 deletion in endothelial senescence and telomerase activity.** Native mesenteric artery endothelial cells were isolated from young (Y) and aged (A) individuals; n=6/group with each sample a pool of 5 different human arteries. (**A**) Representative western blot and respective quantification of Glut1 in young human endothelial cells treated with a control siRNA (siCTL) or an siRNA directed against Glut 1 (siGlut1) for 72 hours. (**B**) Senescence associated β galactosidase (SAβG) staining in young human endothelial cells treated as in A and kept for 4 additional passages in culture. Bar = 20 µm. (**C**) Relative mRNA levels of the senescence marker genes, p21 and p16, in cells as in panel B. (**D**) Telomerase activity measured in young endothelial cells treated as in panel B and additionally receiving ATP-polyamine-biotin for 3 days. (**E**) Immunoprecipitation of telomerase reverse transcriptase (TERT) and immunoblotting for TERT and HSP9a in young cells treated as in panel D. *P<0.05, ***P<0.001. Student's t-test (B,C) or two-way ANOVA (D).

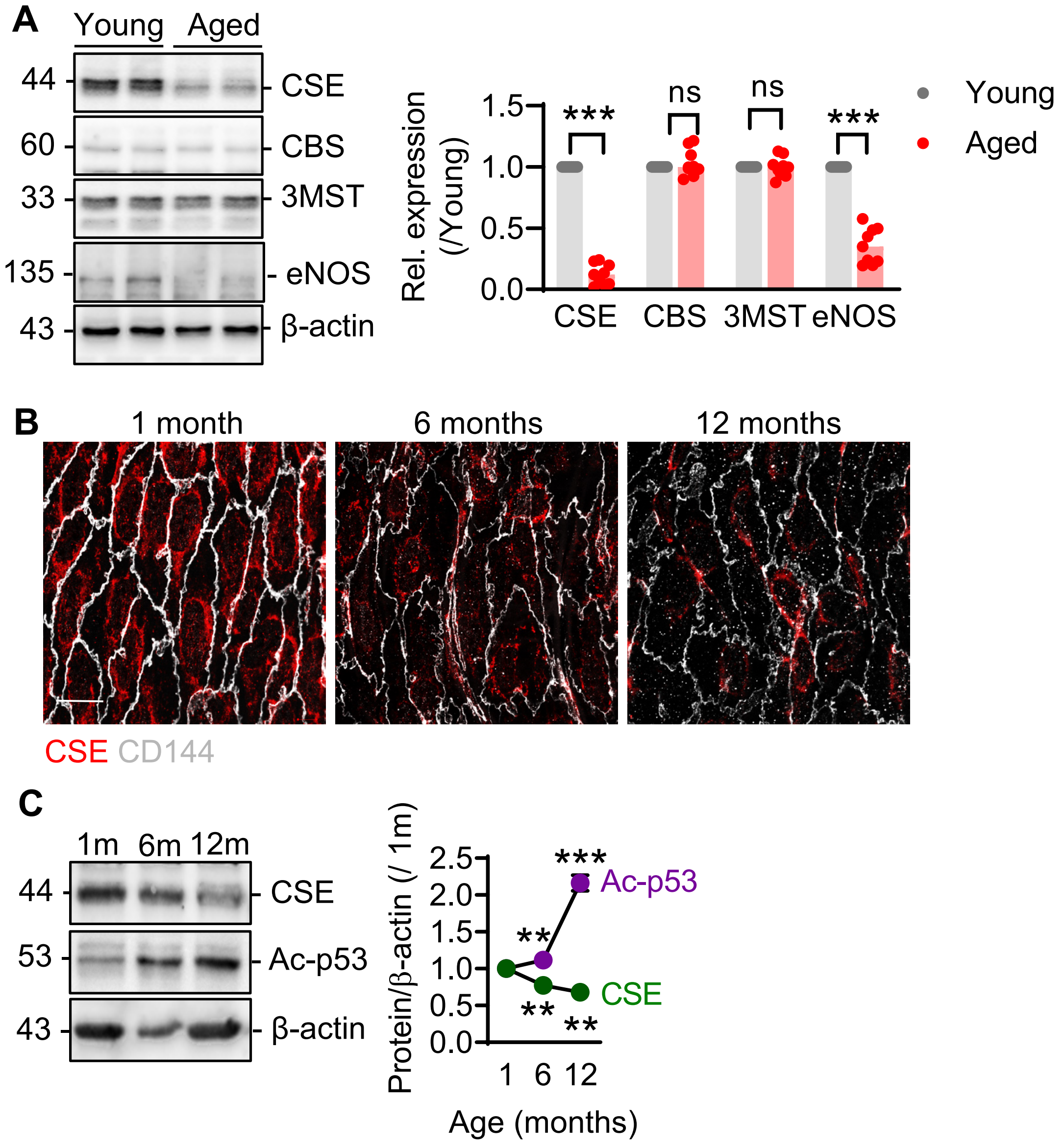


**Supplementary Fig. 4. Effects of senescence in CSE activity and p53 acetylation.** (**A**) Native mesenteric artery endothelial cells were isolated from young (Y) and aged (A) individuals; n=6-9/group with each sample a pool of 5 different human arteries. Immunoblotting and relative quantification of eNOS, CSE, CBS, 3MST and β-actin levels. (**B**) *en face* staining of CSE (red) in CD144+ (grey) endothelial cells from the abdominal aortae from 1, 6, and 12 month old wild-type mice, bar = 10 µm. Comparable results were obtained in 5-7 additional animals. (**C**) CSE expression and p53 acetylation at K120 (Ac-p53) in endothelial cells from 1,6 and 12 month (m) old wild-type mice; n=6/group. **p<0.01, ***p<0.001. Students t-test (A), one-way ANOVA followed by Tukey's multiple comparisons test (C).

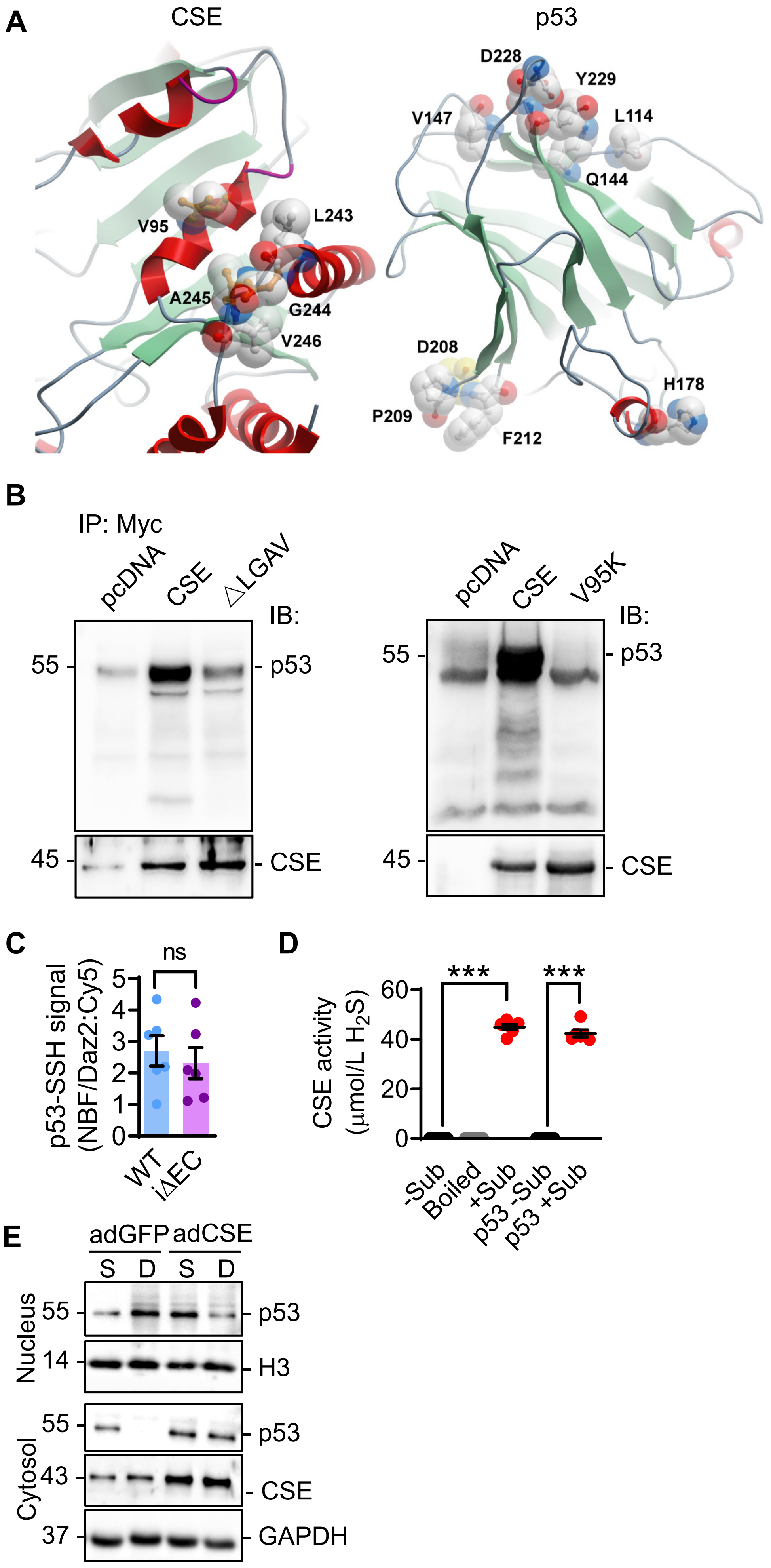


**Supplementary Fig. 5. Interaction between p53 and CSE and its consequences for p53 *S*-sulfhydration and CSE activity.** (**A**) *in silico* detection of the interacting amino acids in CSE and p53. (**B**) Co-immunoprecipitation of pcDNA, wild type human CSE or CSE with a deletion of the LGAV binding sequence (ΔLGAV) or the mutation of Val95 to lysine (V95K) in HEK239 cells**.** n=4/group**.** (**C**) Lack of *S*-sulfhydration of p53 (p53-SSH) detected as the relative fluorescence intensity of NBF/Daz2:Cy5 in endothelial cells from wild-type mice and endothelial cell specific CSE knockout mice (iΔEC); n=6. (**D**) Activity of purified CSE in the absence and presence of p53, and in the absence (-) and presence (+) of the substrates (Sub: L-cysteine, L-cystathionine, both 100 µmol/L, and teh cofactor pyridoxal-5-phosphate, 10 nmol/L); n=6/group. Boiled/denaturated samples were included as a negative control. (**E**) p53, GAPDH and H3 levels in cytoplasmic and nuclear extracts from aged human native endothelial cells. Cells were adenovirally transduced to express either GFP (adGFP), or CSE (adCSE, 10 MOI) 48 hours prior to treatment with solvent (S) or doxorubicin (D, 1 µmol/L, 24 hours); n = 3 different isolates. ***p<0.0001. Student's t test (C), one way ANOVA and Bonferroni multiple comparisons test (D).

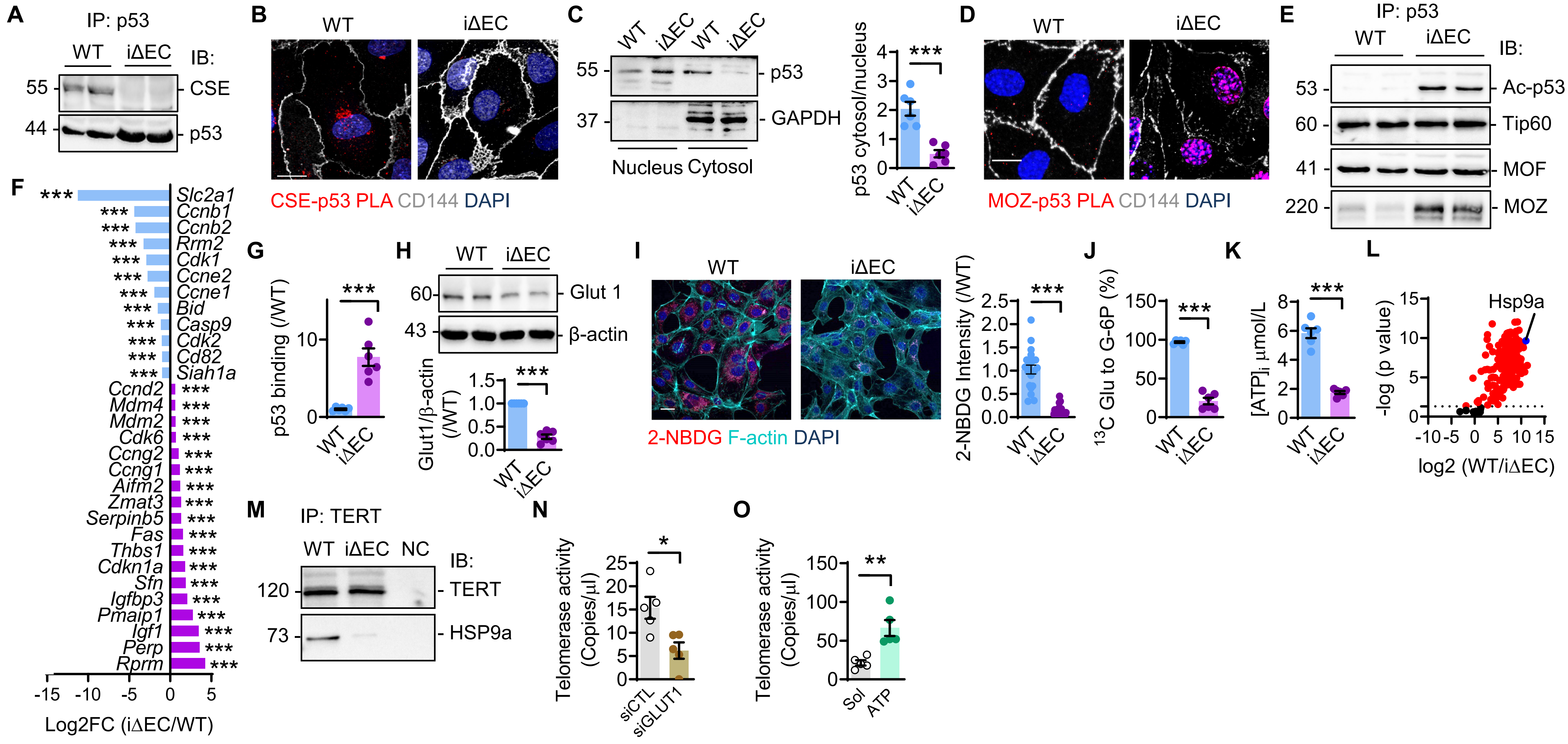


**Supplementary Fig. 6. Consequences of CSE deletion on p53 activation and the telomerase complex.** Pulmonary endothelial cells were isolated from wild-type (WT) and CSE$^{i\Delta EC}$ (iΔEC) littermates. (**A**) Interaction of p53 with CSE in immunoprecipitates (IP). Comparable results were obtained using 6 independent cell batches. (**B**) Proximity ligation assay (PLA) showing the interaction of CSE with p53. Bar = 5 µm. Results represent 5 additional independent experiments. (**C**) Representative immunoblot for p53 and quantification in nuclear and cytosolic extracts. Histone 3 (H3) identifies nuclear fractions; n=6/group. (**D**) Proximity ligation assay (PLA) showing the interaction between MOZ and p53; n=6/group. Bar = 5 µm. (**E**) Representative blots showing the acetylation of p53 (Ac-p53) at K120, and its association with Tip60, MOF and MOZ; n=6/group. (**F**) Heatmap showing the log2 fold change (FC, RNA sequencing) for KEGG pathway 04115-enriched p53 targets; n=4/group. (**G**) p53 binding to the Glut1 promoter identified by Chip-qPCR; n=4/group (each sample is a pool of 2 batches). (**H**) Glut1 expression; n=6/group. (**I**) 2-NBDG (red) uptake. Bar = 5 µm; n=6/ group. (**J**) Contribution (%) of $^{13}$C glucose (Glu) to glucose 6 phosphate (G6P); n=6/group. (**K**) Intracellular concentration of ATP; n=6/group. (**L**) Telomerase complexome identified by LC/MS-MS; n=6/group. (**M**) Interaction of TERT with Hsp9a. NC = negative control (rabbit IgG); (**N-O**) Telomerase activity in wild-type endothelial cells treated with a control siRNA or siRNA directed against Glut1 (N), and in CSE$^{i\Delta EC}$ endothelial cells treated with ATP-polyamine-biotin (O); n=4-5/group. *P<0.05, **P<0.01, ***P<0.001, Student's t-test (C, F, G, H, I, J, K, N, O).

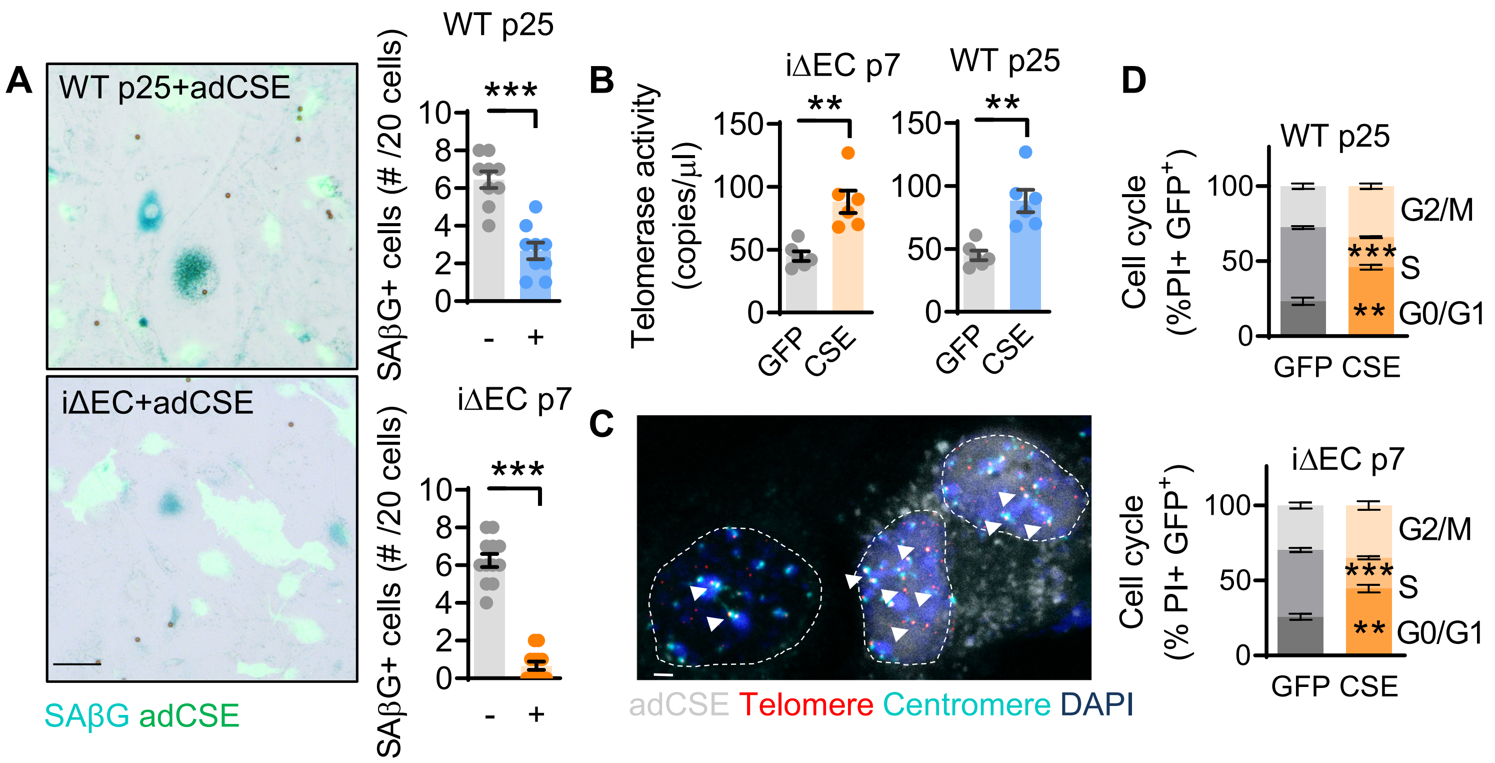


**Supplementary Fig. 7. Effects of CSE overexpression on endothelial senescence.** Murine endothelial cells from wild type (WT) and CSE$^{i\Delta EC}$ mice (iΔEC) were kept in culture and transduced after passage 4 with adenvirsuses encoding GFP or CSE (10 MOI) for up to passage 25 for the wild-type cells, or passage 7 for the CSE$^{i\Delta EC}$ cells. (**A**) Superimposed phase contrast and fluorescent images showing SAβG staining in cells expressing (+) or lacking (-) CSE. Bar = 5 µm; n=12/group. (**B**) Telomerase activity assessed by ddTRAP; n=6/group. (**C**) Telomeres (red) and centromeres (green) identified by PNA-FISH (Bar = 1 µm) in adjacent CSE-deficient and CSE expressing cells. The arrowheads indicate telomeres. Similar results were obtained using 5 additional cell batches. (**D**) Cell cycle analysis in cells from wild-type (WT) mice passaged 25 times or from CSE$^{i\Delta EC}$ (iΔEC) mice passaged 7 times and transduced to express either GFP or CSE; n=6/group. **P<0.01, ***P<0.001. Student's t-test (A, B) or two-way ANOVA followed by Tukey's multiple comparisons test (D).

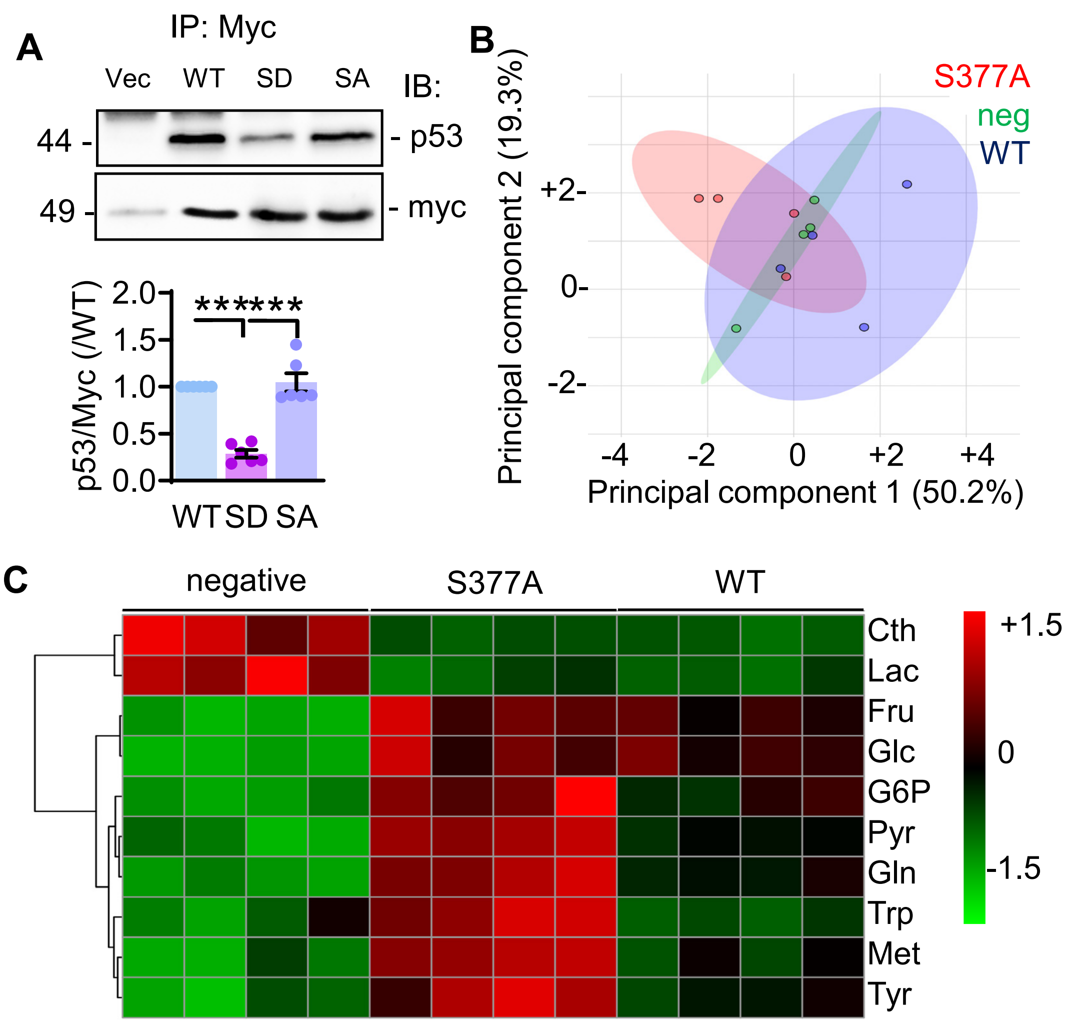


**Supplementary Fig. 8. Effects of CSE mutations on p53 binding and metabolic profile in murine endothelial cell isolated from the aorta**. (**A**) Immunoprecipitation of Myc tagged CSE and western blot for myc (indicative of CSE expression) and p53 in HEK239 cells overexpressing a wild-type (WT) human CSE, or the S377D and S377A mutants. As a negative control cells transfected with the empty vector (Vec) were used. n=4/group. ***P<0.001 (one-way ANOVA followed by Tukey's multiple comparisons test ). (**B-C**) Eight week old $CSE^{i\Delta EC}$ mice were injected AAV9 particles encoding a constitutively active mCherry CSE S377A (S377A) and cells were harvested after 9 months. Negative (neg) stands for CD144+ cells from the same animals that did not express mcherry. Cell isolates were compared with CD144+ endothelial cells isolated from an eight week old wild type (WT) mouse. 2D score plots of the metabolic profile (B). Heatmap of the 10 most altered metabolites (C); n=3-4/group, with each sample a pool of cells from 3-4 animals.
